# Profiling of small non-coding RNAs across cellular and biofluid compartments: implications for multiple sclerosis immunopathology

**DOI:** 10.1101/2020.05.15.097519

**Authors:** Galina Yurevna Zheleznyakova, Eliane Piket, Maria Needhamsen, Michael Hagemann-Jensen, Diana Ekman, Mohsen Khademi, Faiez Al Nimer, Patrick Scicluna, Omid R Faridani, Tomas Olsson, Fredrik Piehl, Maja Jagodic

## Abstract

Multiple sclerosis (MS), a chronic inflammatory disease of the central nervous system (CNS), is associated with dysregulation of microRNAs (miRNA). We here analyzed all classes of small non-coding RNAs (sncRNAs) in matching peripheral blood mononuclear cells (PBMCs), plasma, cerebrospinal fluid (CSF) cells and cell-free CSF from relapsing-remitting (RRMS, n=12 in relapse, n=11 in remission), secondary progressive (SPMS, n=6) MS patients and non-inflammatory and inflammatory neurological disease controls (NINDC, n=11; INDC, n=5). We show widespread changes in small nuclear, nucleolar, transfer RNAs and miRNAs. In CSF cells, 133/133 and 115/117 differentially expressed sncRNAs are increased in RRMS relapse compared to remission and RRMS compared to NINDC, respectively. In contrast, 65/67 differentially expressed PBMC sncRNAs are decreased in RRMS compared to NINDC. The striking contrast between periphery and CNS suggests that sncRNA-mediated mechanisms, including alternative splicing, RNA degradation and mRNA translation, regulate the transcriptome of pathogenic cells primarily in the target organ.

## Introduction

Multiple sclerosis (MS) is a leading cause of neurological disability in young adults, affecting more than 2.3 million people worldwide^1^. The pathology of MS is characterized by the periodic disruption of the blood-brain barrier and infiltration of immune cells, including T cells, B cells and macrophages, into the central nervous system (CNS) resulting in demyelination, neuro-axonal degeneration and neurological deficits. Both genes and environment contribute to disease risk, however, the exact molecular mechanisms that trigger MS are still unclear^2, 3^. Carrying the *HLA-DRB1*15:01* haplotype represents the strongest genetic risk factor, implicating a critical role of CD4^+^ T cells reacting to self-antigens presented by the HLA class II molecules^3, 4^.

Most MS patients (∼85%) initially present with a relapsing-remitting (RRMS) disease course characterized by acute exacerbations followed by periods of full or partial recovery^1^. However, with time more than half will convert to a stage of progressive worsening, i.e. secondary progressive MS (SPMS). MS is diagnosed based mainly on its clinical manifestations, which however are heterogeneous and not unique for MS^1^. While radiological and laboratory tests are therefore increasingly important to provide diagnostic specificity, easily measured biomarkers that can facilitate the diagnostic work up and provide means of monitoring disease activity and treatment response are still largely lacking.

Small non-coding RNAs (sncRNAs) are important regulators of gene expression at the transcriptional and post-transcriptional level and consist of several classes that exert distinct, but overlapping functions^5^. The most functionally investigated sncRNAs are microRNAs (miRNAs), which regulate gene expression by binding to complementary sequences on the 3’ untranslated region of a target messenger RNA (mRNA), leading to translational repression or mRNA degradation^6^. Dysregulation of miRNAs has been described in a range of autoimmune diseases, suggesting involvement in underlying cellular immune mechanisms. Moreover, miRNAs packaged in extracellular vesicles can target gene expression in a recipient cell and thereby modulate immune reactions distally^7^. Additionally, due to their stability in biofluids, miRNAs have been proposed as attractive diagnostic and prognostic biomarkers. Indeed, more than 60 studies have profiled miRNAs among different MS forms and treatment conditions in a variety of biofluids and cellular compartments^8^. The most consistently upregulated miRNAs in MS include miR-142-3p, miR-146a/b and miR-155, while family members of miR-15, miR-548 and let-7 exhibit downregulation^8^. Functionally, these miRNAs have been implicated in immune processes including differentiation and function of CD4^+^ T helper cells^9, 10, 11^. Other miRNAs, such as miR-150 and miR-181c, which are upregulated in MS, have been proposed as biomarkers of different stages of disease, including early active MS^12, 13^. However, studies that profile miRNAs using unbiased genome-wide approaches and investigate other classes of sncRNAs are still scarce^8^.

Downregulation of several small nucleolar RNAs (snoRNAs) has been reported in peripheral blood mononuclear cells (PBMCs), T cells and plasma of MS patients^14, 15, 16, 17^. SnoRNAs are subdivided into the C/D box snoRNAs (SNORDs) that guide methylation of ribosomal RNA (rRNA) and H/ACA box snoRNAs (SNORAs) that assist in pseudouridylation of rRNA. In addition, snoRNAs have been implicated in chromatin remodeling, post-transcriptional genes silencing, splicing and stress responses^18, 19, 20^. Small Cajal body-associated RNAs (scaRNAs) represent a subset of snoRNAs that participate in the maturation of another class of sncRNAs, i.e. small nuclear RNAs (snRNAs) that in the form of ribonucleoprotein (U-snRNPs) complexes are crucial components of the spliceosome. Increased levels of abnormally processed U1, U2, U4, U11 and U12 snRNAs, as well as Y1 RNA, have been described in mononuclear cells of RRMS patients^21^. Y RNAs are components of Ro ribonucleoproteins, which are involved in DNA replication^22^. Recent studies have investigated transfer RNAs (tRNAs) as potential biomarkers in other diseases than MS^23, 24^. Beyond their main role in translation, tRNAs have been shown to be involved in various other processes including cellular homeostasis and gene expression^25^. However, studies investigating these sncRNA classes in the target CNS compartment in MS are still lacking.

Our objective was to perform a comprehensive unbiased sncRNA analysis in matching PBMCs, plasma, cerebrospinal fluid (CSF) cells and cell-free CSF from MS patients and controls. To overcome issues related to the limited RNA input, particularly in CSF cells and cell-free CSF, we used the Small-seq method originally developed for single-cell analysis^26, 27^. Our data demonstrate changes in several classes of sncRNAs, in particular snRNAs, snoRNAs, tRNAs and miRNAs, in MS patients. Moreover, we reveal distinct patterns of sncRNAs across different compartments with important implications for their functional interpretation and assessment of biomarker potential.

## Results

### Detection of sncRNAs

We utilized Small-seq^27^ that incorporates unique molecular identifiers (UMIs) to quantify sncRNA transcripts (Supplementary Fig. 1a). After quality control and pre-processing, sequencing reads were mapped to hg38, amplicons were collapsed and sncRNAs identified based on alignment length of 18-40 nt (Supplementary Fig. 1b). A strict transcript filter of at least 2 UMIs per sample was applied and transcripts were normalized using the trimmed mean of M values (TMM)-method (Supplementary Fig. 1b, Supplementary Fig. 3). Technical replication of three samples from each compartment demonstrated high reproducibility (Fig. 1a).

**Figure 1.**
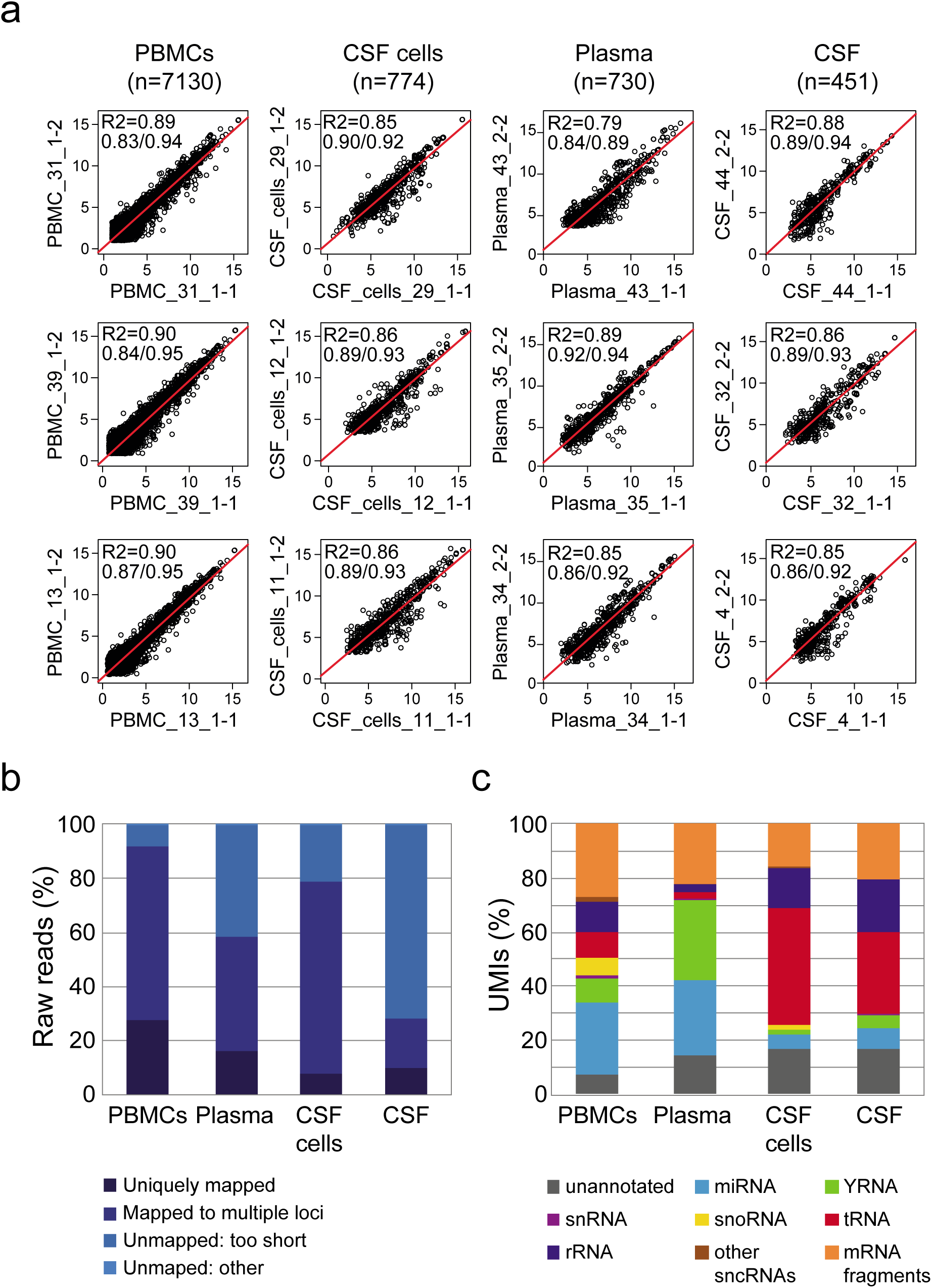
Detection of small non-coding RNAs (sncRNAs). (**a**) Comparison of normalized, log2-transformed counts per million (lcmp) from original samples (X-axis) and technical replicates (Y-axis) from peripheral blood mononuclear cells (PBMCs), cerebrospinal fluid (CSF) cells, plasma and cell-free CSF (n=3 replicates per compartment). The R2, followed by Spearman and Pearson correlation coefficients, are given for each comparison as well as a regression line (red). Included are transcripts with at least 2 unique molecular identifiers (UMIs) in all samples for the compartment of interest, with n referring to the number of transcripts examined (**b**) Distribution of raw reads and (**c**) fraction of UMI counts for different sncRNA classes across each cellular and biofluid compartment. Raw UMI counts across all samples were used to analyze the distribution of different sncRNA classes in each compartment. Classes of snRNAs include microRNA (miRNA) and fragments of ribosomal RNA (rRNA), small nuclear RNA (snRNA), small nucleolar RNA (snoRNA), transfer RNA (tRNA), Y RNA (YRNA), sncRNAs in other classes than the aforementioned (other sncRNAs) and messenger RNA (mRNA) fragments. See also Supplementary Fig. 4, Supplementary Table 1 and Supplementary Data 1.

As expected, the average percentage of aligned reads was higher in the cellular compartments compared to biofluids, with the lowest fraction observed in cell-free CSF (Fig. 1b, Supplementary Table 1, Supplementary Data 1), similar to previous reports^28, 29^.

We classified unique RNA molecules into several categories; miRNAs and fragments of snRNAs, snoRNAs, tRNAs, Y RNAs, rRNAs and mRNAs (Fig. 1c, Supplementary Fig. 4). These fragments can represent full sncRNAs or be fragments actively derived from them. Transcripts with low abundance were collectively included in an “other sncRNAs” category (Supplementary Data 2), while transcripts mapping to the human genome without overlap with any of the RNA categories were classified as “unannotated”. Similar to other studies^29, 30, 31^, we observed a distinct profile of sncRNA classes across analyzed cellular and biofluid compartments (Fig. 1c). While miRNAs were the most abundant class of sncRNAs in PBMCs and plasma (26.4% and 27.9%, respectively), Y RNAs were dominant in plasma (29.6%), consistent with previous reports^30, 32^. On the other hand, tRNAs represented the most abundant class in CSF cells and cell-free CSF (43.6% and 30.3%, respectively) as previously suggested^29^.

Thus, in addition to demonstrating high technical reproducibility over a range of samples with limited RNA input, our method was concordant with previous studies with respect to the classes of detected sncRNAs and their relative abundance.

### Patterns of sncRNAs in MS patients

Next we investigated sncRNAs in PBMCs, plasma, CSF cells and cell-free CSF samples from RRMS (relapse, n=12; remission, n=11) and SPMS (n=6) patients, as well as from non-inflammatory and inflammatory neurological disease controls (NINDC, n=11; INDC, n=5) (Table 1). We first focused on the comparison between RRMS and NINDC (Supplementary Data 2), representing the groups of clinical interest with the largest number of individuals, while relatively small SPMS and INDC groups were only used for a group-level overview (Supplementary Note 1). We then compared the relapse phase of RRMS, characterized by recent worsening of symptoms and/or evidence of inflammatory activity in the CNS detected by imaging, with remission, a phase of stable symptoms without such signs (Supplementary Data 3). An overview on the total number of detected sncRNAs for each class and compartment is summarized in Table 2 and the most abundant transcripts are provided in Supplementary Tables 2-6.

**Table 1.**
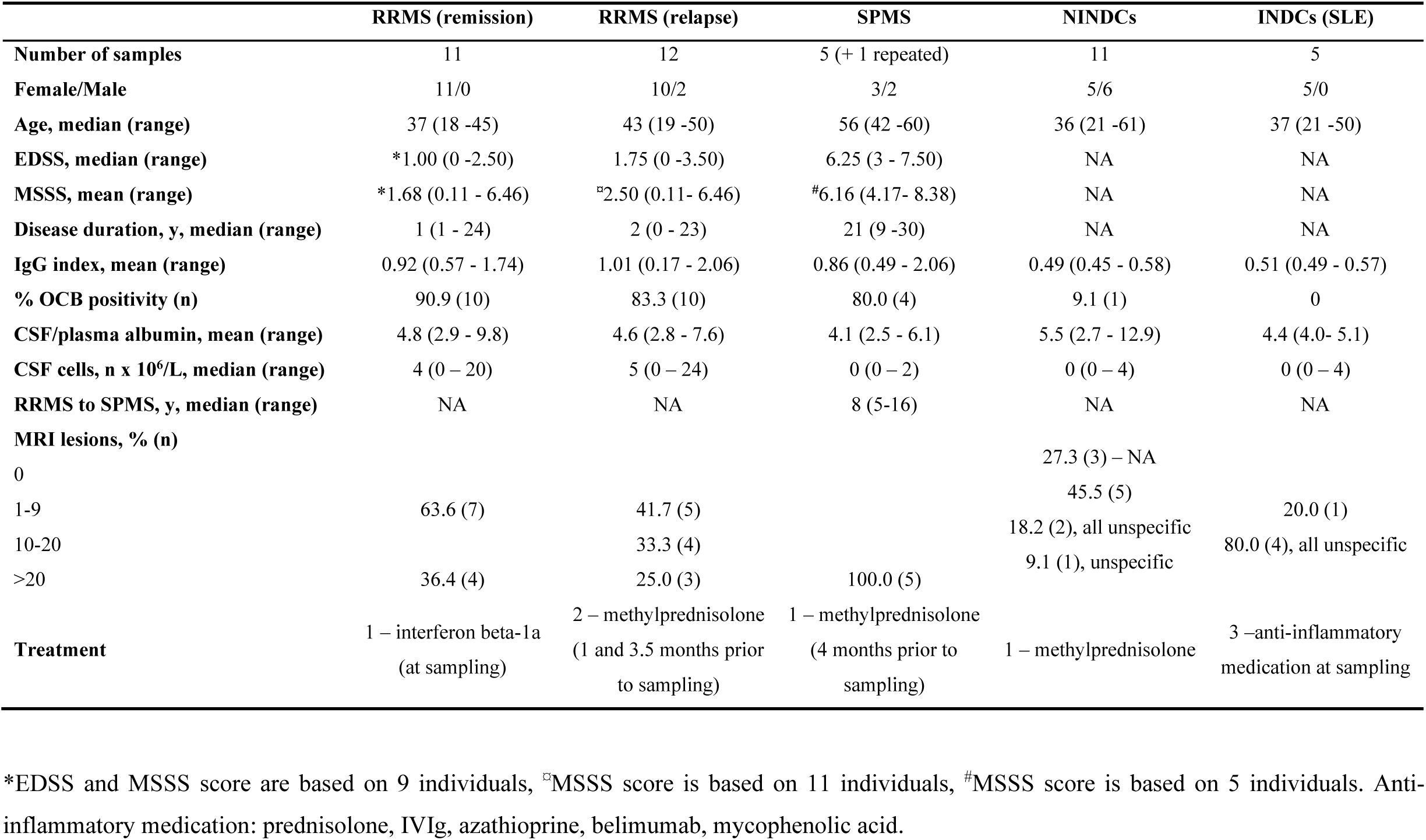
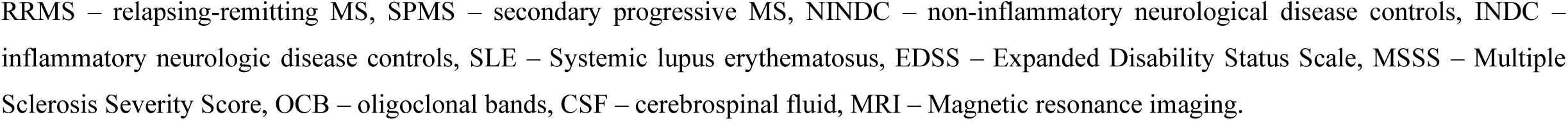
Cohort characteristics.

**Table 2.** Total number of detected small non-coding RNA (sncRNA) fragments in each class across four compartments.

|  | PBMC | Plasma | CSF cells | CSF |
| --- | --- | --- | --- | --- |
| RRMS vs. NINDC analysis |  |  |  |  |
| Total sncRNAs | 1673 | 539 | 469 | 265 |
| miRNAs | 459 | 212 | 91 | 62 |
| tRNAs | 395 | 157 | 200 | 120 |
| snoRNAs | 303 | 8 | 58 | 0 |
| snRNAs | 181 | 25 | 33 | 1 |
| YRNAs | 212 | 82 | 30 | 30 |
| Other sncRNAs | 41 | 13 | 10 | 1 |
| 5S, 5.8S rRNAs | 82 | 42 | 47 | 51 |
| Total all transcripts | 6957 | 671 | 575 | 376 |
| Relapse vs. Remission analysis |  |  |  |  |
| Total sncRNAs | 1706 | 617 | 597 | 334 |
| miRNAs | 475 | 224 | 117 | 82 |
| tRNAs | 395 | 178 | 227 | 159 |
| snoRNAs | 308 | 7 | 93 | 0 |
| snRNAs | 188 | 31 | 44 | 1 |
| YRNAs | 217 | 112 | 42 | 36 |
| Other sncRNAs | 40 | 19 | 16 | 1 |
| 5S, 5.8S rRNAs | 83 | 46 | 58 | 55 |
| Total all transcripts | 8282 | 785 | 753 | 471 |

A total number of detected sncRNAs in each class and total transcripts number counted after the filtering threshold of 2 unique molecular identifiers (UMIs). miRNA – microRNA, tRNA – transfer RNA, snoRNA – small nucleolar RNA, snRNA – small nuclear RNA, YRNA – Y RNA, rRNA – ribosomal RNA.

Notably, while the majority of differentially expressed sncRNAs (adj. p < 0.05) were upregulated in RRMS in the CSF compartment, they were downregulated in PBMCs and plasma compared to NINDC, i.e. displaying an opposing pattern between the two compartments (Fig. 2). In CSF cells, 115/117 differentially expressed sncRNAs were upregulated in RRMS, while in PBMCs 65/67 differentially expressed sncRNAs were downregulated compared to NINDC (Fig. 2a, Supplementary Data 2). Comparison of RRMS relapse and remission identified 133 differentially expressed sncRNAs in CSF cells, all of them upregulated during relapse in RRMS patients, contrasting with a striking pattern of downregulation in PBMCs (Fig. 2a, Supplementary Data 3). Although only one differentially expressed sncRNA was detected in plasma and none reached the significance threshold in cell-free CSF, the patterns generally followed changes observed in the respective cellular compartments (Fig. 2b).

**Figure 2.**
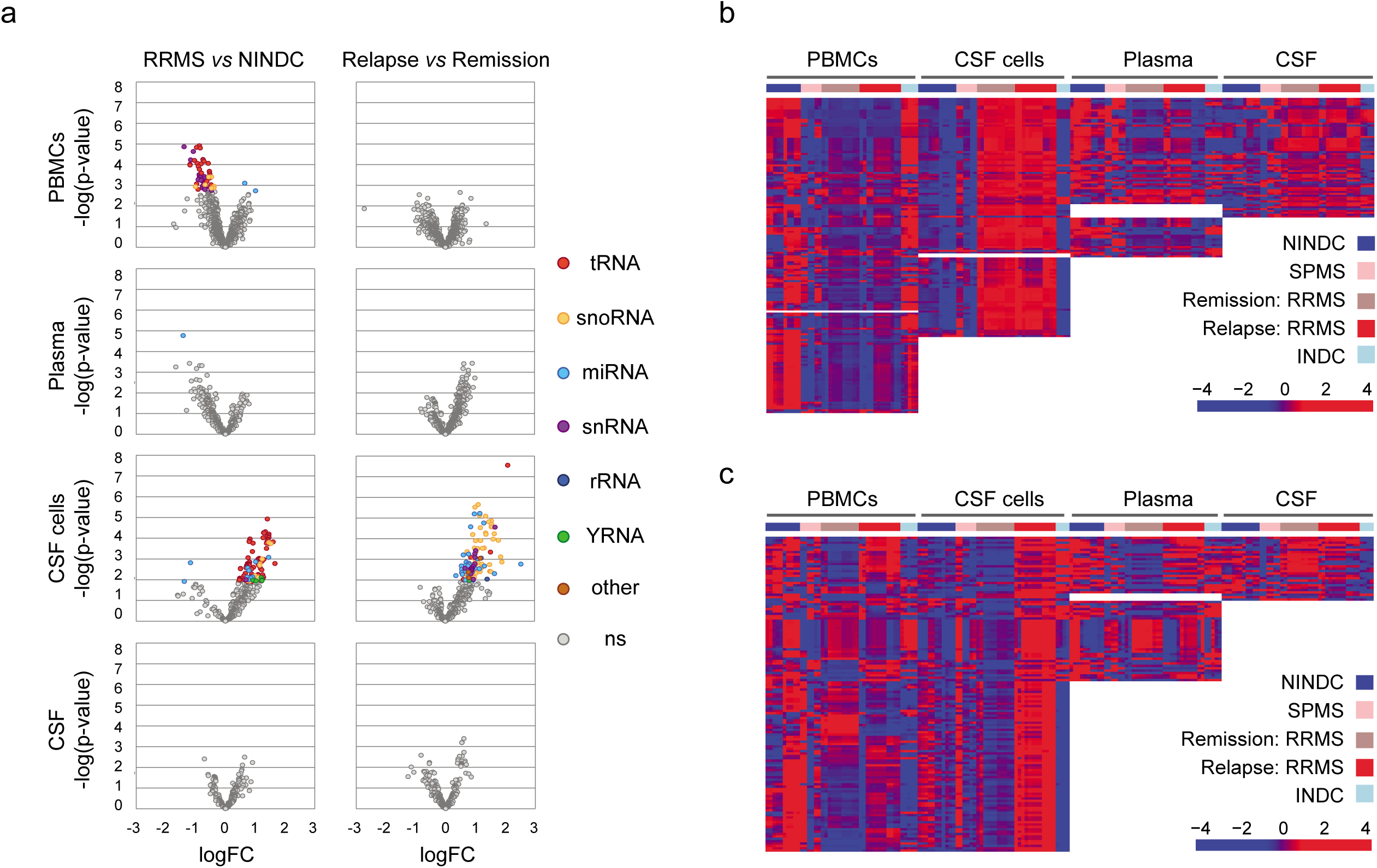
Small non-coding RNAs (sncRNAs) fragments associating with multiple sclerosis (MS) status and phase across four compartments. Small-seq^26, 27^ was utilized to quantify sncRNAs in peripheral blood mononuclear cells (PBMCs), cerebrospinal fluid (CSF) cells, plasma and cell-free CSF from relapsing-remitting MS (RRMS; n=12 in relapse, n=11 in remission), secondary progressive MS (SPMS; n=6), non-inflammatory and inflammatory neurological disease controls (NINDC, n=11; INDC, n=5). (**a**) Volcano plots illustrate differences in several classes of sncRNAs, depicted in different colors, between RRMS vs. NINDC and relapse vs. remission (*Methods*). The y-axis and x-axis depict –log_10_(p-value) and log_2_(fold-change), respectively. Colored circles indicate significant sncRNAs (adj. p-value < 0.05): microRNA (miRNA) and fragments of ribosomal RNA (rRNA), small nuclear RNA (snRNA), small nucleolar RNA (snoRNA), transfer RNA (tRNA), Y RNA (YRNA) and other (other) sncRNAs, ns – not significant. Heatmaps of sncRNAs identified between (**b**) RRMS vs. NINDC and (**c**) relapse vs. remission (adj. p-value < 0.05). Normalized, fitted lcmp values were centered and scaled for each compartment separately, with high levels illustrated in red and low levels in blue. Row-wise hierarchical clustering (Euclidean distance, Complete) was conducted based on RRMS-derived transcripts in PBMCs or CSF cells after sub-clustering NAs (white block), representing transcripts that did not pass the filtering threshold. Column-wise hierarchical clustering (Euclidean distance, Complete) was conducted separately within each compartment and patient group.

These observations suggest distinct patterns of sncRNAs across different compartments in MS patients. In the following sections, we present a detailed investigation of the most frequently differentially expressed sncRNAs. While we report only significant sncRNAs (adj. p < 0.05) in the following text, all sncRNAs with nominal p < 0.01 were included in heatmaps to investigate sncRNA patterns across patient groups and compartments (Fig. 3-5).

**Figure 3.**
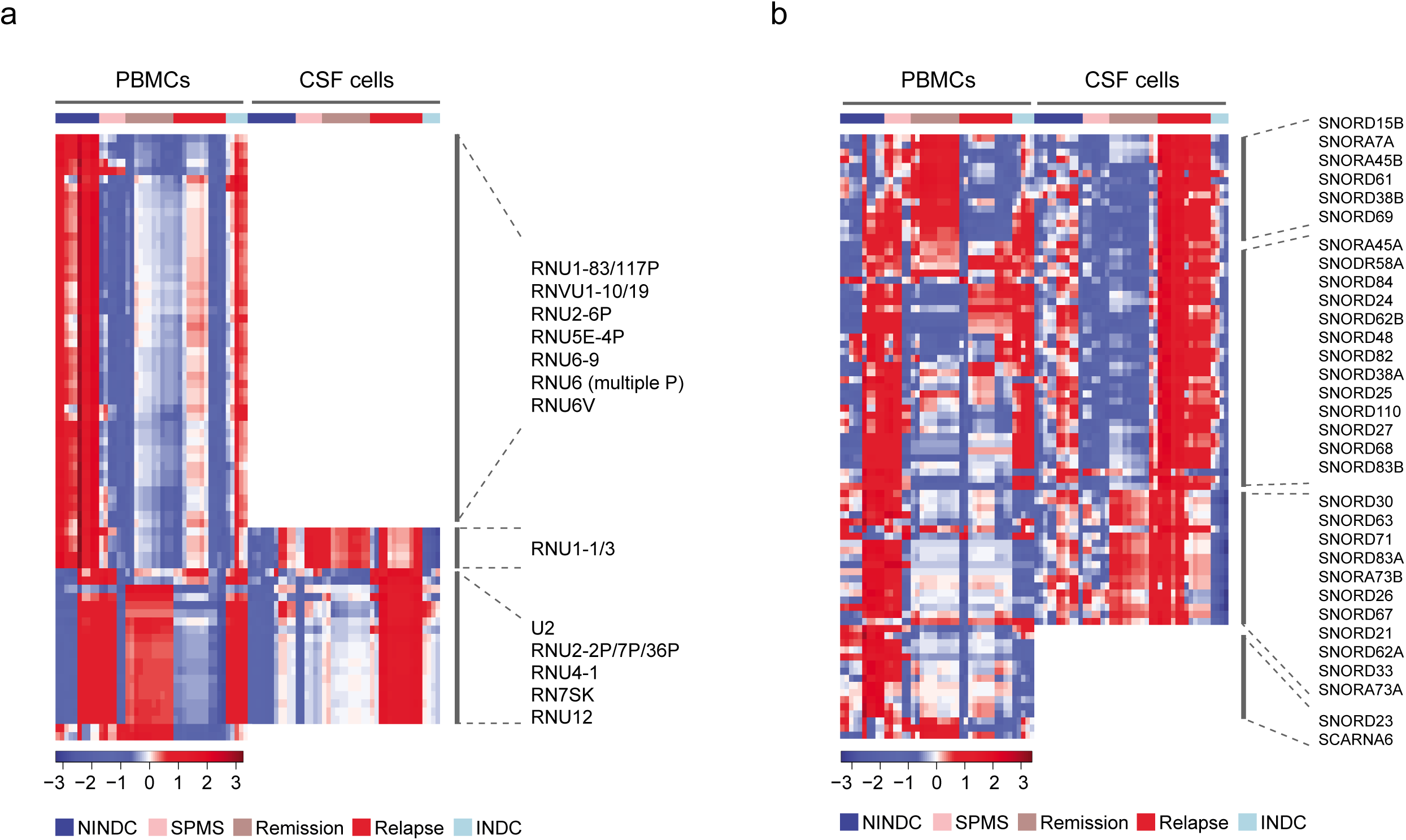
Small nuclear (snRNA) and nucleolar (snoRNA) RNA fragments associating with multiple sclerosis (MS) status and phase across intracellular compartments. Heatmaps of (**a**) snRNAs and (**b**) snoRNAs identified using Small-seq^26, 27^ in peripheral blood mononuclear cells (PBMCs) and cerebrospinal fluid (CSF) cells from relapsing-remitting MS (RRMS; n=12 in relapse, n=11 in remission), secondary progressive MS (SPMS; n=6), non-inflammatory and inflammatory neurological disease controls (NINDC, n=11; INDC, n=5). Transcripts identified between RRMS vs. NINDC and/or relapse vs. remission (p-value < 0.01) were included. Distinct groups of snRNAs and snoRNAs, depicted by the vertical lines and representative molecules to the right of each heatmap, were differentiated based on their profile in PBMCs and CSF cells. Both heatmaps contain normalized, fitted lcmp values, which were centered and scaled within the PBMC and CSF cell compartment separately with high levels illustrated in red, low levels in blue and intermediate levels in white (see color key). Row-wise hierarchical clustering (Euclidean distance, Complete) was conducted based on PBMC and/or CSF cell, RRMS-derived transcripts after sub-clustering NAs (white block), which represent transcripts that did not pass the filtering threshold. Column-wise hierarchical clustering (Euclidean distance, Complete) was conducted separately within each compartment and patient group.

**Figure 4.**
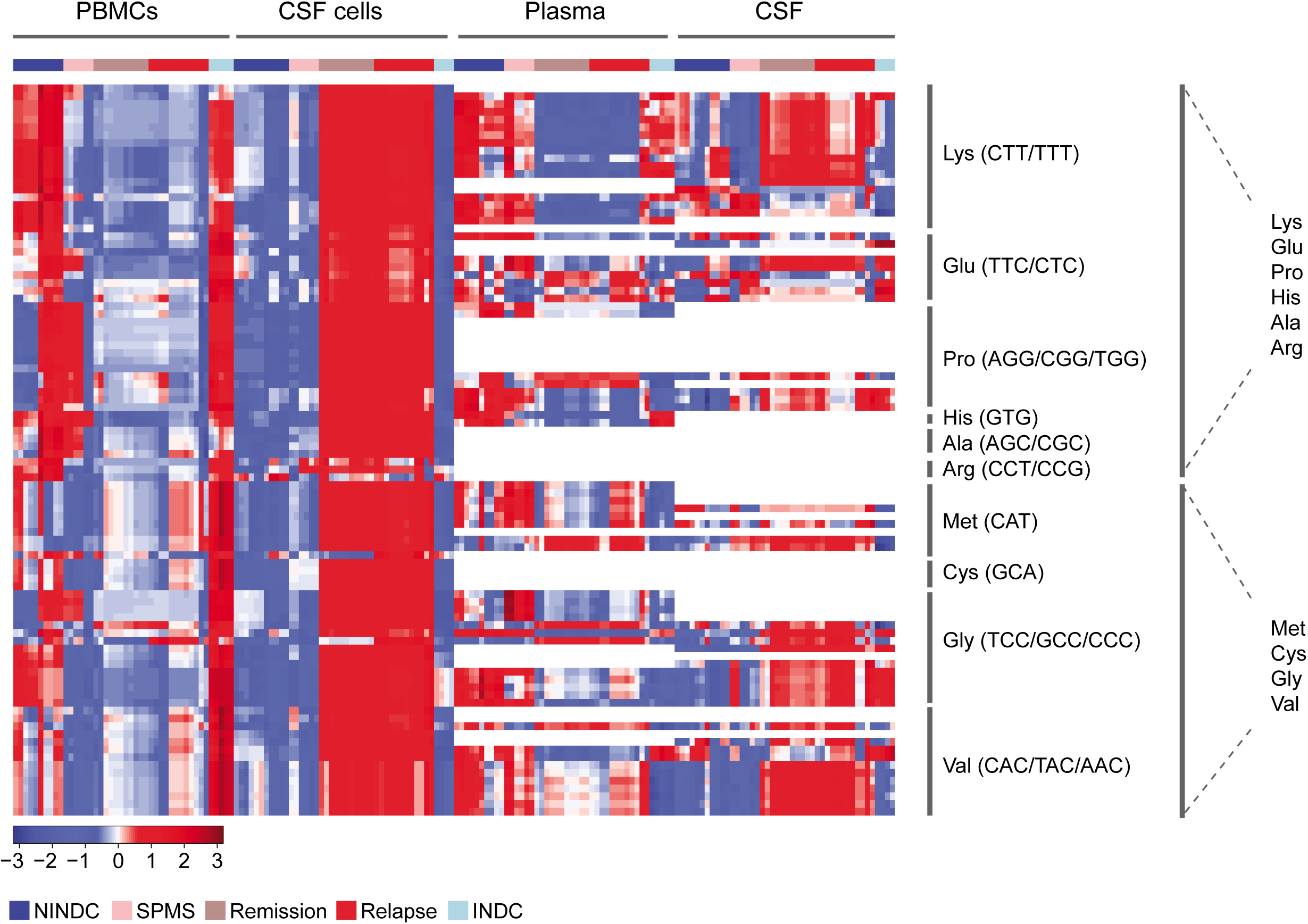
Transfer RNA (tRNA) fragments associating with multiple sclerosis (MS) status and phase across four compartments. (**a**) Heatmap of tRNAs identified using Small-seq^26, 27^ in peripheral blood mononuclear cells (PBMCs), plasma, cerebrospinal fluid (CSF) cells and cell-free CSF from relapsing-remitting MS (RRMS; n=12 in relapse, n=11 in remission), secondary progressive MS (SPMS; n=6), non-inflammatory and inflammatory neurological disease controls (NINDC, n=11; INDC, n=5). Selected transcripts identified between RRMS vs. NINDC and/or relapse vs. remission (p-value < 0.01) in PBMCs and CSF cells were included. tRNAs were grouped based on amino acid origin, depicted by vertical lines with representative amino acid and tRNA anticodon(s) to the right of the heatmap. Two distinct groups of tRNAs, depicted by second vertical lines and representative amino acid to the right of the heatmap, were differentiated based on the tRNA profile in PBMCs and CSF cells. The heatmap contains normalized, fitted lcmp values, which were centered and scaled within each compartment separately with high levels represented in red, low levels in blue and intermediate levels in white (see color key). Row-wise hierarchical clustering (Euclidean distance, Complete) was based on PBMC and CSF cell compartments after sub-clustering based on amino acid origin. NAs (white block) represent transcripts that did not pass the filtering threshold. Column-wise hierarchical clustering (Euclidean distance, Complete) was conducted separately within each compartment and patient group.

**Figure 5.**
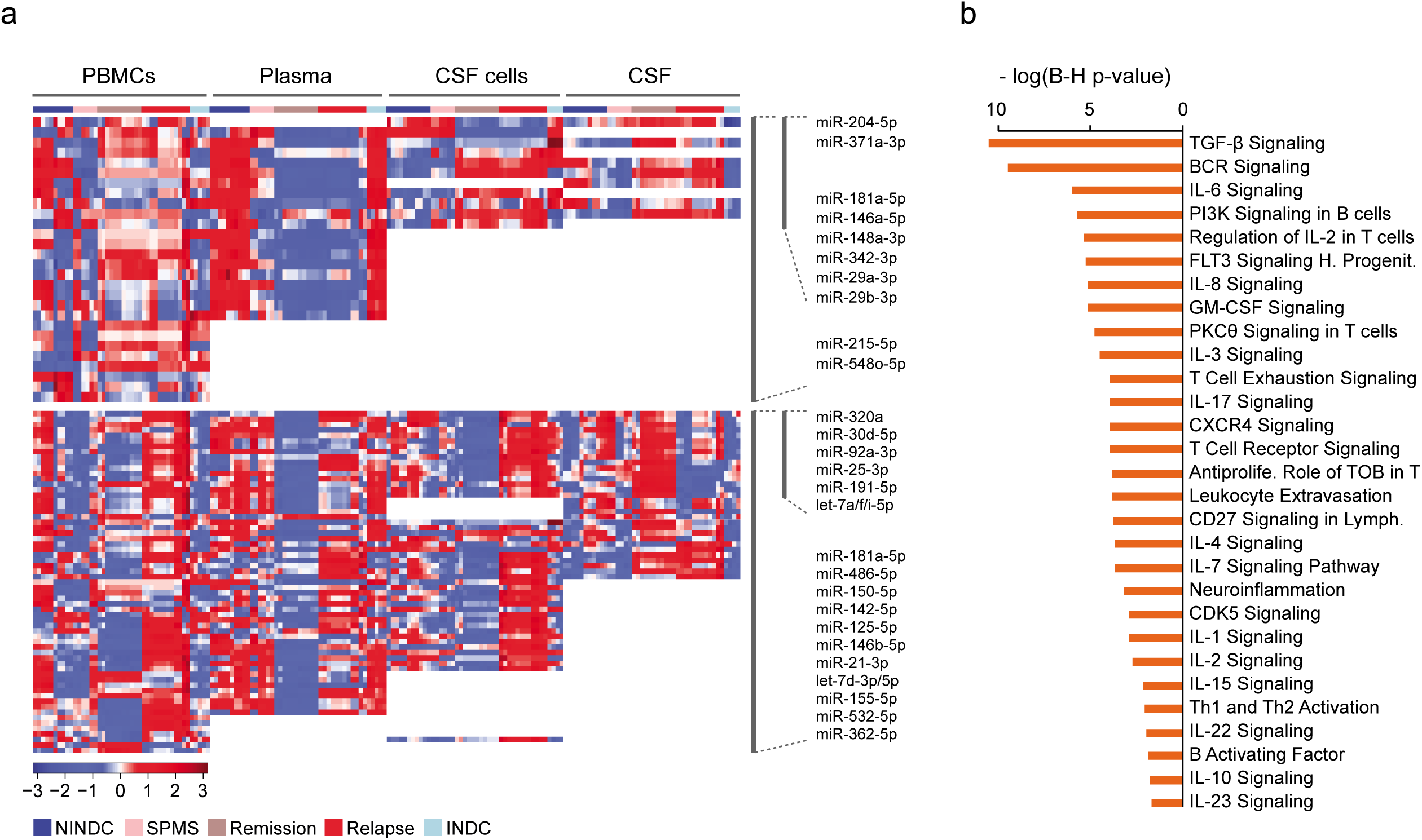
MicroRNAs (miRNA) associating with multiple sclerosis (MS) status and phase across four compartments. (**a**) Heatmap of miRNAs identified using Small-seq^26, 27^ in peripheral blood mononuclear cells (PBMCs), plasma, cerebrospinal fluid (CSF) cells and cell-free CSF from relapsing-remitting MS (RRMS; n=12 in relapse, n=11 in remission), secondary progressive MS (SPMS; n=6), non-inflammatory and inflammatory neurological disease controls (NINDC, n=11; INDC, n=5). Transcripts identified between RRMS vs. NINDC and/or relapse vs. remission (p-value < 0.01) were included. Two distinct groups of miRNAs, depicted by the vertical lines and representative molecules to the right of the heatmap, were differentiated based on the miRNA profile across the compartments. The heatmap contains normalized, fitted lcmp values, which were centered and scaled within the compartments separately with relatively high levels illustrated in red, relatively low levels in blue and intermediate levels in white (see color key). Row-wise hierarchical clustering (Euclidean distance, Complete) was conducted based on CSF, CSF cell or PBMC, RRMS-derived transcripts (relapse vs. remission) after sub-clustering based on NAs (white block), representing transcripts that did not pass the filtering threshold. Column-wise hierarchical clustering (Euclidean distance, Complete) was conducted separately within each compartment and patient group. (**b**) Significant immune-related canonical pathways (Benjamini-Hochberg-corrected p–value < 0.05) generated using Ingenuity Pathway Analysis on predicted target genes of miRNAs differentially expressed between relapse vs. remission in CSF cells.

#### snRNA profile

We identified fragments of canonical snRNAs as well as “variant” snRNAs, which are encoded by pseudogenes of multiple snRNAs genes^33^. Differentially expressed snRNAs were detected only in the cellular compartments, likely due to their nuclear localization and thus low abundance in biofluids (Fig. 1c). In total, we identified 33 differentially expressed snRNAs (adj. p < 0.05) in PBMCs between RRMS and NINDC, while in CSF cells, 19 snRNAs were differentially expressed (adj. p < 0.05) between RRMS relapse and remission (Fig. 2a, Supplementary Data 2-3).

Three distinct groups were differentiated based on their patterns of expression across the compartments (Fig. 3a). Members of the largest group comprised 31 snRNAs (adj. p < 0.05) that were detected only in PBMCs and displayed downregulation in RRMS compared to NINDC (Fig. 3a, Supplementary Data 2). This group included canonical and variant/pseudogene snRNA transcripts of U1, U6 clusters and U11 transcript. A second, smaller group, comprising two U1 snRNAs (adj. p < 0.05), was also downregulated in PBMCs of RRMS, but displayed a clear opposing pattern of upregulation in CSF cells compared to NINDC (Fig. 3a). The third and last group comprised 19 snRNAs (adj. p < 0.05) that were upregulated during RRMS relapse compared to remission in CSF cells, but tended to be downregulated in PBMCs (Fig. 3a, Supplementary Data 3). Both canonical and pseudogene transcripts of the U2-1 cluster, U4 and U12 as well as the RN7SK transcript belonged to this group.

Thus, while U1/U6 transcripts were predominantly detected in PBMCs and showed downregulation in RRMS compared to NINDC, U2 transcripts were more specific for disease activity with upregulation in CSF cells and downregulation in PBMCs in relapse compared to remission.

#### snoRNA profile

Similar to snRNAs, due to their nucleolar localization and low abundance in biofluids, we identified snoRNA fragments predominantly in cellular compartments (Fig. 1c). A total of seven and ten differentially expressed snoRNAs (adj. p < 0.05) were detected between RRMS and NINDC in PBMCs and CSF cells, respectively. We also identified 65 differentially expressed snoRNAs between RRMS relapse and remission in CSF cells (Fig. 2a, Supplementary Data 2-3).

The differential expression profile in PBMCs and CSF cells separated snoRNAs into four distinct groups (Fig. 3b). The first group included 14 snoRNAs (adj. p < 0.05) that were upregulated in RRMS relapse in CSF cells, while displaying an opposing pattern of being downregulated in PBMCs compared to remission (Fig. 3b, Supplementary Data 3). The group contained snoRNAs that predominately participate in modifications of 28S rRNA (Supplementary Table 4) and is represented by six H/ACA (including SNORA7A) and eight C/D box snoRNAs (such as SNORD69). Similar to the pattern in the first group, members of the second group, comprising 34 snoRNAs (adj. p < 0.05), were upregulated in RRMS relapse compared to remission in CSF cells, but in contrast to the first group displayed no clear differences in PBMCs (Fig. 3b). This group consisted predominantly of C/D box snoRNAs, including the *U22* locus snoRNAs (SNORD25 and SNORD27).

The third group comprised 17 snoRNAs (adj. p < 0.05) that were upregulated in RRMS relapse compared to remission in CSF cells, similar to the first two groups (Fig. 3b, Supplementary Data 3). They all also showed a clear pattern of upregulation in RRMS compared to NINDC in CSF cells, with 9/17 being significantly different (adj. p < 0.05). However, they displayed an opposing pattern in PBMCs reflecting predominant downregulation in RRMS compared to NINDC, with 5/17 displaying significant difference (adj. p < 0.05). This group comprised C/D box snoRNAs, including another two *U22* locus snoRNAs (SNORD26 and SNORD30) and two H/ACA box snoRNAs (SNORA73A and SNORA73B). The last fourth group was only expressed in PBMCs, not CSF cells, and exhibited the same pattern as the previous group, including C/D box snoRNA SNORD23 and SCARNA6 (adj. p < 0.05) that were downregulated in RRMS compared to NINDC (Fig. 3b, Supplementary Data 2).

Thus, the vast majority of differentially expressed snoRNAs were upregulated in CSF cells from RRMS patients, specifically during the relapse phase, while the same snoRNAs were frequently downregulated compared to NINDC controls in PBMCs.

#### tRNA profile

We detected 25 and 91 differentially expressed tRNA fragments (adj. p < 0.05) between RRMS and NINDC in PBMCs and CSF cells, respectively (Fig. 2a, Supplementary Data 2). In total, ten tRNAs displayed differential expression (adj. p < 0.05) between RRMS relapse and remission in CSF cells (Fig. 2a, Supplementary Data 3). For further analysis, we grouped tRNAs based on their amino acid origin (Supplementary Fig. 5) and separated them into two main groups based on their differential expression patterns across the compartments (Fig. 4).

The first group of tRNAs, coding for Lys, Glu, Pro, His, Ala and Arg, was characterized by strong dysregulation between RRMS and NINDC and a contrasting pattern between CSF cells and PBMCs (Fig. 4). A total of 19 Lys tRNAs (both TTT and CTT isoacceptors) were upregulated in RRMS (adj. p < 0.05) in CSF cells, most of them (17/19) displaying downregulation compared to NINDC in PBMCs (Fig. 4, Supplementary Data 2). In addition, ten Glu (both CTC and TTC isoacceptors), 11 Pro (all isoacceptors), three His (GTG) and four Ala (TGC, CGC, AGC) tRNAs were elevated in RRMS compared to NINDC (adj. p < 0.05) in CSF cells. In contrast, the vast majority of aforementioned tRNAs displayed a clear trend for downregulation in RRMS compared to NINDC in PBMCs (Fig. 4, Supplementary Data 2). Several of the Lys, Glu and Pro tRNAs were also detected in cell-free CSF and plasma, where their pattern mirrored the pattern in the corresponding cellular compartments, i.e. CSF cells and PBMCs (Fig. 4, Supplementary Data 2).

The second group of tRNAs, coding for Met, Cys, Gly and Val, displayed significant differential expression only in CSF cells (Fig. 4, Supplementary Data 2). The levels of nine Met tRNAs (involved in both initiation and elongation) were significantly higher in RRMS compared to NINDC (adj. p < 0.05). Additionally, five Cys, 13 Gly (GCC, TCC, CCC isoacceptors) and 16 Val (all isoacceptors) tRNAs also displayed the same pattern of upregulation in RRMS compared to NINDC in CSF cells (adj. p < 0.05). Several of the Gly and Val tRNAs were also detected in cell-free CSF, where their pattern mirrored the pattern observed in CSF cells (Fig. 4, Supplementary Data 2).

Taken together, differentially expressed tRNAs were particularly enriched in CSF cells, and displayed extensive upregulation in RRMS patients compared to controls, which contrasted with their unchanged or downregulated profile in PBMCs.

#### miRNA profile

A total of 13 differentially expressed miRNAs (adj. p < 0.05) were detected between RRMS and NINDC, ten in CSF cells, two in PBMCs, and one in plasma (Fig. 2a, Supplementary Data 2). In addition, 33 miRNAs were differentially expressed (adj. p < 0.05) between relapse and remission in CSF cells of RRMS patients (Fig. 2a, Supplementary Data 3).

Based on the miRNA expression profile across all compartments, we differentiated two main groups of miRNAs (Fig. 5a). Members of the first group of miRNAs were differentially expressed between RRMS and NINDC across multiple compartments (Fig. 5a, Supplementary Data 2). A set of ten miRNAs within this group also differed in CSF cells (adj. p < 0.05), with miR-146a-5p, miR-148a-3p, miR-150-5p, miR-181a-5p, miR-29a/b-3p and miR-342-3p being upregulated, while miR-204-5p and miR-371a-3p were downregulated in RRMS compared to NINDC. In PBMCs, two miRNAs including miR-548o-3p were significantly upregulated (adj. p < 0.05) in RRMS compared to NINDC with many other miRNAs showing the same trend (Fig. 5a). In contrast, plasma miRNAs displayed predominant downregulation, with miR-215-5p being significantly downregulated (adj. p < 0.05) in RRMS compared to NINDC (Fig. 5a, Supplementary Data 2).

Members of the second group were predominantly upregulated in the relapse phase in PBMCs, plasma and, in particular, in CSF cells where 33 miRNAs were significantly upregulated in relapse compared to remission (adj. p < 0.05) (Fig. 5a, Supplementary Data 3). This group included miR-125a-5p, miR-146a/b-5p, miR-150-5p, miR-155-5p, miR-181a-5p, miR-21-3p and miR-320a, together with several members of the let-7 family, such as let-7a/d/f/i-5p. Interestingly, a subset of them, including miR-21-5p, miR-92a-3p and let-7 members, displayed an opposing pattern in cell-free CSF and CSF cells between relapse and remission (Fig. 5a, Supplementary Data 3).

To explore the possible functional implications, we performed Ingenuity Pathway Analysis (IPA) on the predicted targets of the detected miRNAs. We focused on most frequently differentially expressed miRNAs, i.e. the 33 miRNAs in CSF cells that were upregulated in RRMS relapse compared to remission. Many of the identified (adj. p < 0.05) immune-related pathways concerned activation of T and B cells, as well as cytokine and chemokine signaling, with transforming growth factor beta (TGF-β) signaling being the most significantly enriched pathway (Fig. 5b).

The miRNA profile thus underscores a general upregulation of miRNAs in RRMS, particularly in CSF cells during the relapse phase, with possible implications for regulation of T and B cell activation and differentiation.

#### Other small-noncoding RNAs

We also observed changes in the expression level of other sncRNAs (Fig. 2a, Supplementary Data 2). For example, significantly higher levels of signal recognition particle RNA transcript (RN7SL1) as well as RNA component of mitochondrial RNA processing enzyme complex (RNase MRP RNA) and mitochondrially encoded tRNA-Met (MT-TM) were found in CSF cells during RRMS relapse compared to remission (adj. p < 0.05, Supplementary Data 3).

#### mRNA fragments profile

In addition to sncRNAs, we detected four and 28 differentially expressed mRNA fragments (adj. p < 0.05) in PBMCs and CSF cells, respectively, between RRMS and NINDC (Supplementary Data 4), and ten (adj. p < 0.05) in CSF cells between RRMS relapse and remission (Supplementary Data 5). As the nature of such mRNA fragments is still largely unexplored, the outcome of mRNA fragment analysis should be taken with a caution.

To address potential biological functions, we performed IPA on genes that associated with differentially expressed mRNA fragments. While analysis in CSF cells, likely due to the limited number of input genes, yielded no significant pathways, PBMCs displayed sufficient number of genes for IPA analysis (Fig. 6). Four canonical pathways remained significant after Benjamini-Hochberg correction between RRMS and NINDC in PBMCs, including *Eukaryotic Initiation Factor 2* (*EIF2), Signaling* and *Mammalian Target of Rapamycin* (*mTOR) Signaling* (Fig. 6a), with significant enrichment of biological functions connected to *Cell death & Survival* and *Protein Synthesis* (Fig. 6b). Multiple pathways and biological functions were significantly enriched between RRMS relapse and remission in PBMCs including pathways involved in leukocyte activation and migration (Fig. 6a) along with biological functions related to the number and proliferation of lymphocytes (Fig. 6b).

**Figure 6.**
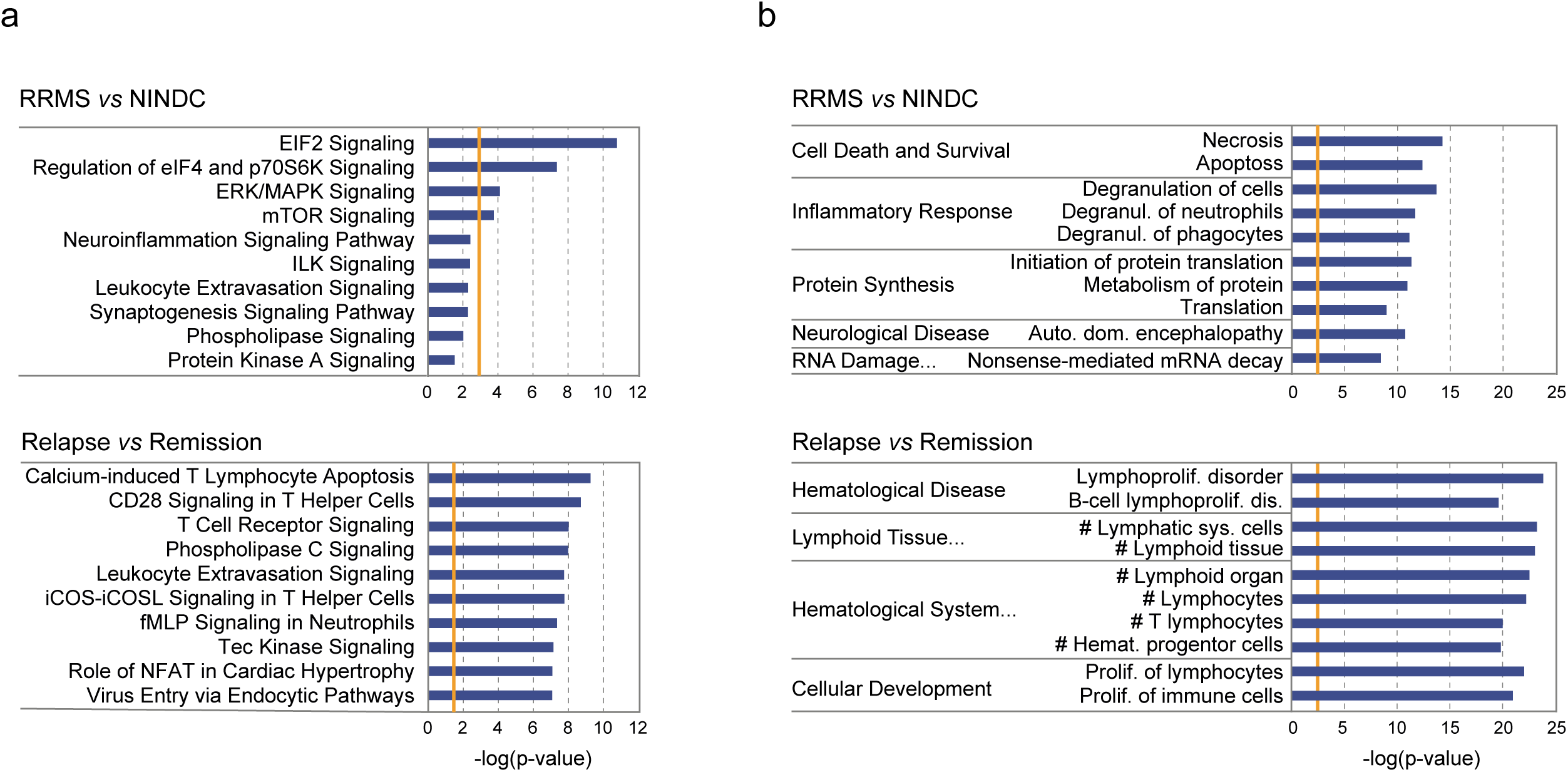
Pathways and functions implicated by messenger RNA (mRNA) fragments associating with multiple sclerosis (MS) status and phase in peripheral blood monocular cells (PBMCs). mRNA fragments were identified using Small-seq^26, 27^ in PBMCs from relapsing-remitting MS (RRMS; n=12 in relapse, n=11 in remission), secondary progressive MS (SPMS; n=6), non-inflammatory and inflammatory neurological disease controls (NINDC, n=11; INDC, n=5). Transcripts associating with candidate differentially expressed mRNA fragments (p-value < 0.1) between RRMS vs. NINDC as well as relapse vs. remission were analyzed using Ingenuity Pathway Analysis (IPA) and top ten (non-cancer) (**a**) Canonical pathways and (**b**) Diseases and functions, comprising minimum ten molecules, are depicted in the figure. The –log_10_ of the Fisher’s p-value is provided on the X-axis, while the Benjamini-Hochberg-corrected threshold for significance is indicated by the orange line. All tissues and confidence levels were used in the IPA analysis.

We next sought to explore the putative contribution of miRNA-mediated regulation of the identified mRNAs and therefore asked whether predicted targets of significantly differentially expressed miRNAs are enriched among the differentially expressed mRNA fragments in the corresponding compartment and comparison. There was no significant enrichment of targets of the differentially expressed miRNAs among mRNA fragments detected in PBMCs and CSF cells from RRMS compared to NINDC (Supplementary Table 7). On the contrary, comparison of RRMS relapse to remission in CSF cells revealed a significant enrichment (Fisher’s test p < 0.05) of miRNA targets, for 30 out of 33 differentially expressed miRNAs, among the dysregulated mRNA fragments (Supplementary Table 7).

Thus, the differentially expressed mRNA fragments suggest alteration of functional pathways in MS patients in general and provide additional evidence in support of the observed miRNA changes during the relapse phase in particular.

## Discussion

We performed a comprehensive sncRNA analysis in MS patients and controls utilizing Small-seq^26, 27^. Our main findings demonstrate widespread alterations of several classes of sncRNAs, particularly during the relapse phase in CSF cells. Furthermore, we report the opposing pattern of snRNA, snoRNA and tRNA changes between the blood and CNS compartments. The patterns across several sncRNA species implicate changes in the general cellular mechanisms, such as alternative splicing and mRNA translation, occurring for example during lymphocyte activation, while cytokine signaling pathways critical for T helper differentiation were more selectively modulated by miRNAs. Collectively, these observations underscore the relevance of studying the CSF compartment in a CNS disease such as MS, with CSF reflecting more closely disease processes occurring in the target organ^34^, including enrichment of encephalitogenic immune cell populations^35, 36^.

We detected predominant upregulation of snRNAs in CSF cells and downregulation in PBMCs from RRMS patients compared to controls. We also observed increased levels of multiple U1, U2, U4 and U12 snRNAs transcripts in CSF cells during the relapse phase. An altered level of U2-associated Protein SR140 has previously been detected in CSF cells of MS patients compared to NINDC^37^. Similar to our observations in PBMCs, diminished global levels of U1, U5 and U6 transcripts have been reported in PBMCs of RRMS^37^. The observed changes are consistent with alteration of several scaRNAs and snoRNAs that guide posttranscriptional modifications of U2, U5 and U6 snRNAs (Supplementary Table 4). Since snRNAs are crucial regulators of alternative splicing, this pattern strongly suggests disturbances in mRNA splicing, which has already been noted in MS^38^. Indeed, activation of both T and B cells initiates global alternative splicing events in multiple genes of the NF-κB, mitogen-activated protein kinases (MAPK) and Rho GTPase signaling and cell proliferation pathways^39^. Several splicing factors were found altered in MS^40^, including factors that participate in the assembly of the aforementioned snRNAs in spliceosome units. Accordingly, our analysis of mRNA fragments also revealed decreased levels of transcripts deriving from two splicing factors, *SNRPB* and *RBFOX2*, in PBMCs of RRMS patients (Supplementary Data 4). Altered RNA splicing in MS is consistent with increased levels of the long non-coding RNA *MALAT1*, affecting the expression of splicing factors *HNRNPF* and *HNRNPH1*, described in blood of RRMS patients^41^. Interestingly, in our study, while *MALAT1*-associated small cytoplasmic RNA (mascRNA) was downregulated in PBMCs of RRMS patients, levels of mascRNA were augmented in CSF cells in RRMS compared to controls (Supplementary Data 4).

Alterations in alternative splicing mechanisms are further supported by the observed changes of snoRNAs, which exert non-canonical functions in mRNA processing^43,44^. Most of the snoRNAs exhibited downregulation in PBMCs of RRMS compared to controls, which is in agreement with previous studies reporting predominant downregulation of snoRNAs in CD3^+^ T cells and PBMCs of MS patients^8^. Notably, similar to the snRNA profile, these snoRNAs demonstrated the opposite pattern in CSF cells compared to PBMCs, with upregulation in CSF cells in RRMS compared to controls, particularly during the relapse phase. Such pattern was observed for several *U22* locus snoRNAs, including SNORD27 previously shown to regulate alternative splicing and to activate silent exons^42^ thereby potentially contributing to the splicing defects implicated in MS. Notably, the majority of snoRNAs are encoded inside the introns of protein-coding genes and their expression may be altered as a consequence of frequent intron retention due to splicing impairment. Finally, altered alternative splicing is also concordant with changes in miRNAs that can target alternative splicing factors. For example, miR-181a-5p found upregulated in CSF cells of RRMS patients, targets a member of the serine/arginine (SR)-rich family of splicing factor SRSF7^43^. Overall, we show accumulating evidence from different sncRNA classes for aberrant alternative splicing mechanisms in MS.

Other dysregulated snoRNAs exert their canonical function in assisting in rRNA modifications that are essential for accurate ribosome function. Elevated levels of misprocessed 18S and 28S rRNAs have previously been reported in RRMS^21^. The majority of detected snoRNAs are encoded inside structural ribosomal proteins, elongation factors and translational regulators (Supplementary Table 4)^44, 45^ controlled by the mTOR pathway^46^. Since the transcription of such genes is activated when intensive protein production is required, the detected snoRNA alterations may reflect global changes in expression of translational machinery genes. Accordingly, our analysis of differential mRNA fragments implicates *EIF2 Signaling* as the top enriched canonical pathway (Fig. 6) together with changes in many ribosomal structural proteins, translational initiation and elongation factors (Supplementary Data 4) in PBMCs of RRMS compared to controls.

Global changes in translation in RRMS patients are further supported by strong alterations of tRNAs. Most of Lys, Glu, Pro, His and Ala tRNA were found upregulated in CSF cells from RRMS compared to controls. Additionally, Met, Cys, Gly and Val tRNAs displayed robust upregulation in CSF cells in RRMS as well as during the relapse phase. Similar to snRNAs and snoRNAs, tRNA changes in PBMCs exhibited the opposite direction, i.e. the majority of these tRNAs displayed downregulation in PBMCs of RRMS patients. The RNA polymerase III transcribes tRNAs, as well as both U6 snRNAs and RN7SL1, which showed a similar direction of dysregulation in MS patients as tRNAs. This provides additional evidence in support of our hypothesis that changes in tRNA levels might be induced by general changes in tRNA synthesis and processing. In that respect, elevated levels of specific tRNAs in CSF cells may reflect adaptation to specific inflammatory conditions^47, 48^ that initiate changes in protein profile synthesis and abundancy.

The changes in tRNAs, snoRNAs and snRNAs may be linked to changes in mTOR activity, known to facilitate cellular growth and proliferation. Indeed, in our study, the analysis of mRNA fragments in PBMCs implicated mTOR among the most significant canonical pathways. One of the mTOR complexes, mTORC1, promotes mRNA translation and elongation through ribosomal protein S6 kinase 1 (S6K1) and eukaryotic translation initiation factor 4E (eIF4E)-binding protein 1 (4E-BP1) as well as by increasing rRNA and tRNA transcription^49^. Some evidence also suggests the involvement of the mTORC1 complex in U-snRNP biogenesis^50^. Taken together, the striking upregulation of all of the above classes of sncRNAs in CSF cells, particularly in the relapse phase, is consistent with activation of mTOR signaling in activated and proliferating cells enriched in the target organ. In line with this, antigen recognition triggers mTOR signaling via the PI3K-Akt pathway and directs differentiation of naïve T cells into T_H_1, T_H_2 and T_H_17 cells, but inhibits differentiation of T_REG_ cells^51^, by transferring signals from growth and immunomodulatory factors to lineage-selective STAT proteins and transcription factors^51, 52^. Increased levels of leptin, a hormone that activates mTOR signaling^53^, has been reported in the CSF of RRMS patients, where leptin levels correlated with a reduced numbers of T_REGS_ ^54^. Additionally, the hyporesponsiveness of T from RRMS patients *in vitro* and *in vivo* is associated with activation of the leptin-mTOR metabolic pathway^55^.

It is difficult to speculate to what degree the changes we observe here are explained by changes occurring in encephalitogenic cells that are enriched in the CSF compartment and what is due to regulatory mechanisms in the larger population of non-encephalitogenic cells mainly located in the blood compartment. Another level of complexity relates to the temporal dynamics in the regulation of immune cells during the course of MS. For example, autoproliferation of *bona fide* encephalitogenic T and B cell clones is downregulated in blood from MS patients in relapse compared to remission^56^. Nevertheless, the identified changes in components of snRNA-mediated splicing machinery in PBMCs further substantiate previously described alternative splicing disturbances in peripheral blood of MS patients and collectively concur to general disturbances of these processes in immune cells of MS patients^38^ (Fig. 7). Conversely, changes in snoRNA- and tRNA-mediated mRNA translation mechanisms were more conspicuous in CSF cells. In MS, CSF cells preferentially include CCR7-expressing memory CD4^+^ T cells characterized by upregulation of genes involved in T cell activation as opposed to downregulation of genes associated with naïve cell state^36, 37, 57^. Our observations are in accordance with a recent single cell study comparing CSF cells and blood cells of MS patients during relapse^36^. In blood cells they demonstrated an increased transcriptional diversity compared to CSF cells, reflected by an increased proportion of differentially expressed genes across different cell clusters, while CSF cells show an upregulation of cell cycle genes such as CCNC and Cyclin-C^36^. In addition, most of altered snRNAs, snoRNAs and tRNAs had a tendency of being decreased in PBMCs but increased in CSF cells of RRMS compared to INDC, which comprised systemic lupus erythematosus (SLE) patients (Fig. 3-4, Supplementary Note 1). Therefore, one can speculate that while in RRMS patients there is an enrichment of activated proliferating cells with activated mTOR in the CSF compartment (Fig. 7), in SLE patients they may be directed to other organs.

**Figure 7.**
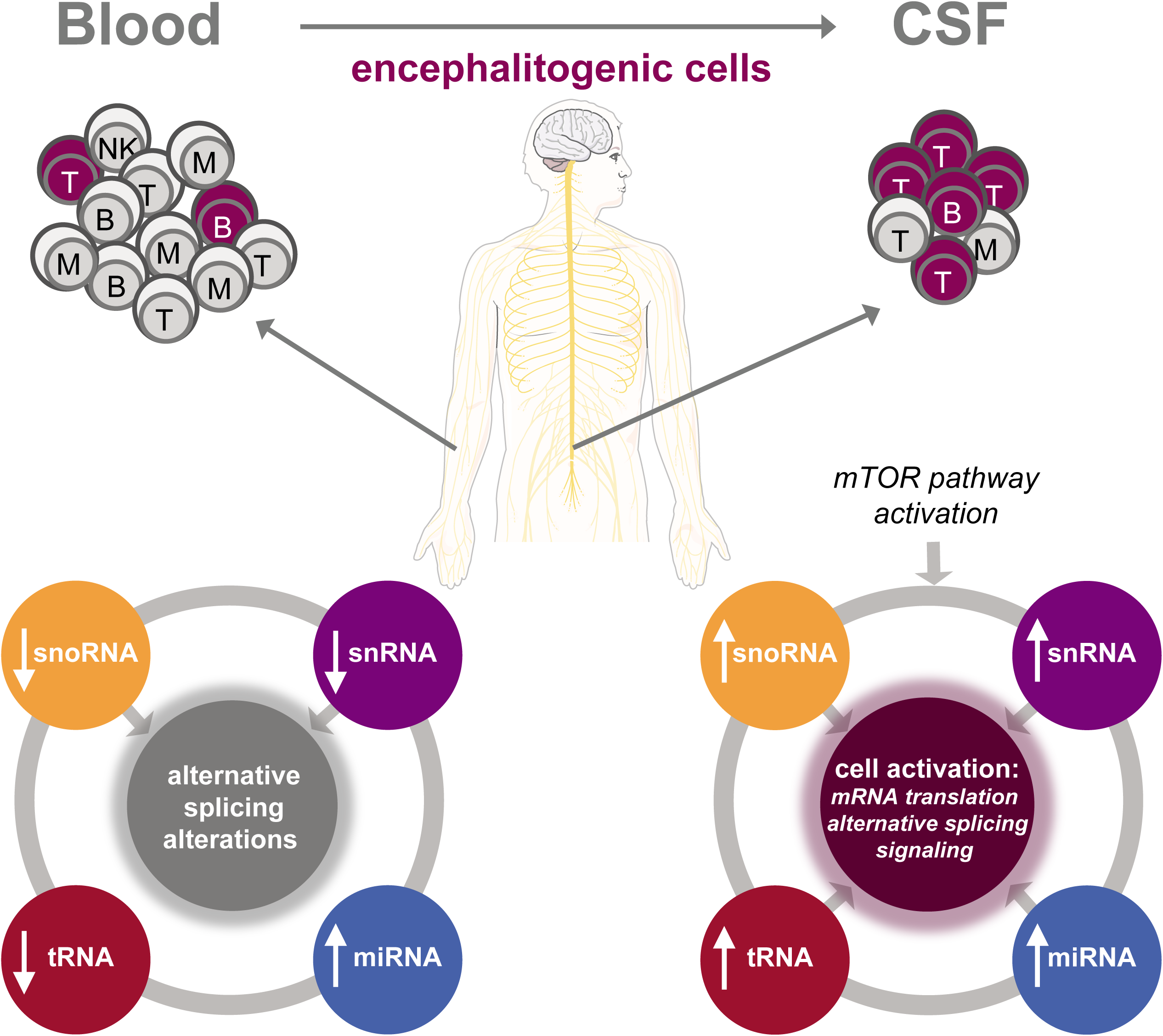
Schematic overview of potential mechanisms of small non-coding RNAs (sncRNAs) in the immunopathology of multiple sclerosis (MS). Most frequently differentially expressed classes of sncRNAs identified in peripheral blood mononuclear and cerebrospinal fluid (CSF) cells in this study include microRNAs (miRNA) and fragments of small nuclear RNAs (snRNA), small nucleolar RNAs (snoRNA) and transfer RNAs (tRNA), with white arrows indicating the direction of a change (increased, decreased) in MS patients. Cellular mechanisms implicated by differentially expressed sncRNAs in blood and CSF cells, the latter being enriched in encephalitogenic cells, are outlined in middle circles and support from a specific class of sncRNA is indicated by grey arrows.

In contrast to most sncRNAs, miRNAs did not display divergent changes between periphery and CNS. This observation supports the notion that inflammation triggers expression of miRNAs in multiple cell types, although the outcome of this upregulation will strongly depend on the target mRNAs content of the host cell. Interestingly, although miRNAs have been the most thoroughly studied sncRNA class to date, differentially expressed miRNAs constitute only a fraction of the altered sncRNAs in MS. Most of the differentially expressed miRNAs were detected in CSF cells during RRMS relapses, with a similar pattern across other compartments. Many of these miRNAs have previously been found to be upregulated in peripheral immune cells from MS patients, i.e. PBMCs and blood cells (e.g. miR-125a-5p, miR-146a-5p, miR-155-5p, miR-362-5p, let-7d-5p) and CD4^+^ T cells (miR-155-5p, let-7i-5p, miR-486-5p)^8^. Additionally, several miRNAs upregulated in CSF cells in relapse compared to remission have also been found dysregulated in the CNS tissue of MS patients; in inactive lesions (miR-155-5p, miR-30d-5p, miR-532-5p), active lesions (miR-146b-5p, miR-320a, miR-142-5p, miR-21-3p), demyelinated hippocampi (miR-30d-5p, let-7f-5p) and cell-free CSF (miR-21-3p, miR-191-5p)^8^. Altogether, these findings suggest that infiltrating immune cells are likely the main cellular source of these CNS-detected miRNAs, although resident CNS cells might be an additional source. Interestingly, IPA analysis of the predicted targets of differentially expressed miRNA between relapse and remission unveiled TGF-β signaling as the most significant pathway. This is in line with previous studies suggesting miRNA-mediated alterations of TGF-β signaling, implicated in differentiation of both T_REG_ and T_H_17 lineages, in MS^8, 58^. Multiple detected miRNAs have been shown to target TGF-β signaling, including relapse-associated miR-21-3p, known to target RBPMS^59^ which regulates TGF-β signaling by increasing transcriptional activity of Smad2/3^60^. Another example is miR-92a, also upregulated in CSF cells during RRMS relapses, which targets SP1^61^ and has been shown to interact with Smad2/3/4^62^. Together with previous studies^8, 58^, our findings position TGF-β signaling as a pivotal mechanism in the immunopathogenesis of MS.

In conclusion, our study demonstrates prominent cellular changes in several classes of sncRNAs with substantial changes correlating with disease status (RRMS vs. NINDC) and disease activity (relapse vs. remission), likely reflecting enrichment of activated encephalitogenic cells in the target organ. Contrasting differentially regulated sncRNA species between the blood and CSF compartments and between relapse and remission highlights the importance of sncRNA-mediated mechanisms, in particular alternative splicing, mRNA degradation and translation, in shaping-up the transcriptome and function of pathogenic cells in MS.

## Methods

### Patients

In total, 23 RRMS and five (one with repeated measurement) SPMS patients, as well as five inflammatory (INDCs) and 11 non-inflammatory neurological disease controls (NINDCs), were included in the study. All MS patients were diagnosed according to the 2010 revised McDonald’s criteria. Patients’ Expanded Disability Status Scale (EDSS) and the Multiple Sclerosis Severity Score (MSSS) were determined by the treating neurologist. The detailed cohort characteristics are presented in Table 1. The study was approved by the Regional Ethical Board, Stockholm, Sweden (ethics permit number: 2009/2107-31/2) and all patients signed the informed consent.

### Sample preparation

#### Preparation of cell-free CSF and CSF cells

The CSF samples were collected in 2 x 15 ml size tubes and were centrifuged immediately after a lumbar puncture at 440 g for 10 minutes at RT to separate cells and larger particles from the CSF supernatants. The CSF supernatants were batched and stored at −80°C and the remaining CSF cells were pooled and concentrated to a volume of 20-60 μl and the cells were transferred to a 2 ml size polypropylene tube, frozen immediately on dry-ice and stored at −80°C until use. One INDC control was missing a CSF cell and cell-free CSF sample.

#### Plasma preparation

The blood samples were collected in EDTA tubes and centrifuged at 1500 g for 15 minutes at RT. The plasma phase was batched and stored at −80°C until use.

#### Preparation of peripheral blood cells

The blood samples were collected in cell preparation tubes (CPT, BD Biosciences) and centrifuged at 1500 g for 15 minutes at RT. Mononuclear cells and platelets form a layer just under the plasma layer. The plasma layer was aspirated, and the cell layer was gently transferred into a 15 ml size tube, 15 ml PBS was added to wash the cells by centrifuging at 300 g for 15 minutes at RT. The cell pellet was re-suspended in 3 ml PBS and distributed into 3 x 2 ml size polypropylene tubes. The cells were centrifuged at 300 g for 10 minutes at RT. The supernatants were aspirated in a way that only cell pellets remained in the tubes. The cell pellets were frozen immediately on dry-ice and stored at −80°C until use. One NINDC control was missing a PBMC and plasma sample.

### RNA extraction

#### RNA isolation from cell-free CSF and plasma samples

sncRNAs were isolated from 300 μl of plasma or CSF using the miRCURY RNA isolation kit for biofluids (Exiqon, Denmark) according to the manufacturer’s protocol. Before isolation CSF and plasma were centrifuged at 3000 g for 5 minutes to remove debris. On-column DNase digestion was performed. RNA was eluted with 15 μl and 20 μl of RNase-free water for CSF and plasma samples, respectively. Eluted RNA was aliquoted and immediately stored at −80°C.

#### RNA isolation from CSF cells and PBMCs

CSF cell precipitates (approximately 5000 cells) was resuspended in 700 μl of QIAzol Lysis Reagent (Qiagen, Germany) and stored at −80°C. After thawing, 200 μl of QIAzol Lysis Reagent and 180 μl of Chloroform (Merck, Germany) was added to the CSF samples. Approximately 1 million PBMCs were used for RNA isolation. After thawing, 900 μl of QIAzol Lysis Reagent and 180 μl of Chloroform was added to the PBMCs samples. Total RNA isolation procedure then continued using the miRNeasy micro kit (Qiagen, Germany) according to the manufacturer’s protocol. On-column DNase digestion was performed. RNA was eluted with 15 μl and 40 μl of RNase-free water for CSF cells and PBMCs samples, respectively. Eluted RNA was aliquoted and immediately stored at −80°C. The quality and quantity of RNA were assessed using small RNA electrophoresis with the 2100 Bioanalyzer System (Agilent, USA) and Qubit RNA high-sensitivity assay (Life Technologies, USA). The concentration range was 100–1500 pg/μl and 0.4–34.6 ng/μl for CSF cells and PBMCs, respectively. Equal RNA amount per compartment was used for preparation of libraries.

### Library preparation and sequencing

#### sncRNA library preparation and sequencing

5 pmol of 5.8S rRNA masking oligo was added to 3 μl of RNA from CSF cells and PBMCs. 0.3 μl of lysis buffer (0.13% Triton-X-100, 4 units of recombinant RNase Inhibitor, Takara) was added to 3.7 μl of cell-free CSF and plasma. The samples were incubated at 72°C for 20 minutes. Further stages were identical for all types of samples and followed the described protocol^26, 27^. Libraries for all 45 samples from each compartment (PBMCs, plasma, CSF cells and cell-free CSF) were prepared simultaneously and were then pooled for each compartment and purified with DNA Clean & Concentrator-5 (ZymoResearch, Germany). The samples were barcoded, using multiplex indexes, to facilitate sequencing of all libraries in each pool on one flow cell lane. Automatic size selection (sizing 125–160 bp) was performed for each library pool using the Pippin Prep (Sage Science, Inc.) to remove primer dimers and other undesirable fragments. The quality and quantity of each library pool were determined on the High Sensitivity DNA electrophoresis with the 2100 Bioanalyzer System (Agilent, USA) and with Qubit dsDNA high-sensitivity assay (Life Technologies, USA). Clustering was done by ‘cBot’ and each pool was sequenced on two lanes of HiSeq2500 (HiSeq Control Software 2.2.58/RTA 1.18.64) with a 1×51 setup using ‘HiSeq SBS Kit v4’ chemistry at the National Genomics Infrastructure (NGI) Stockholm. The Bcl to FastQ conversion was performed using bcl2fastq_v2.19.1.403 from the CASAVA software suite. Phred scores remained above 30 from 1 to 50 bp for all samples (Supplementary Fig. 2a).

#### Technical replication

To replicate our data, PBMCs, plasma, CSF cells and cell-free CSF samples (n=3 per compartment) were chosen from the same cohort. Repeated sncRNA isolation was performed from 300 μl of biofluid samples as described above. A new aliquot of already extracted RNA was used for CSF cells and PBMC samples. The new libraries for all samples were prepared following the above-described protocol. All 12 libraries were pooled together and purified with DNA Clean & Concentrator-5 (ZymoResearch). Automatic size selection (sizing 125–160 bp) was performed using the Pippin Prep (Sage Science, Inc.). The quality and quantity of the pool were determined on the High Sensitivity DNA electrophoresis with the 2100 Bioanalyzer System (Agilent, USA) and with Qubit dsDNA high-sensitivity assay (Life Technologies, USA). Clustering was done by ‘cBot’ and the pool was sequenced on one lane of HiSeq2500 Rapid mode (HiSeq Control Software 2.2.58/RTA 1.18.64) with a 1×101 setup using ‘HiSeq Rapid SBS Kit v2’ chemistry at the National Genomics Infrastructure (NGI) Stockholm. The Bcl to FastQ conversion was performed using bcl2fastq_v2.20.0.422 from the CASAVA software suite. Phred scores remained above 30 from 1 to 74 bp for all samples (Supplementary Fig. 2b).

### Data analysis

#### Preprocessing and read alignment

Preprocessing and alignment of reads was done according to Hagemann-Jensen *et al.*^27^. In short, the UMI sequences were removed from the FastQ files and appended to its corresponding header. Adapter sequences and CA bases were removed with CutAdapt (v1.8.1) and trimmed reads were mapped to hg38 using STAR (v2.4.0). PCR amplicons were then collapsed using the UMI reads. From these, RNA molecule counts, reads aligning from 18 to 40 nt were appointed. Subsequently, transcripts were annotated using Gencode (v22), miRbase (v21) and GtRNAdb (v1) databases.

#### Sequencing statistics

All samples passed the quality control and were included in the downstream analysis. The detailed sequencing statistic including total number of input reads, alignment parameters, total number of UMIs for each compartment is summarized in Supplementary Table 1 and the detailed statistic about each sample is provided in Supplementary Data 1.

#### Transcript filter

Transcripts with less than 2 UMIs in more than 80% of individual samples per contrasted group (i.e. RRMS, NINDC, relapse and remission) were filtered out. Transcripts derived from sncRNAs were identified using the getBM() function from the Bioconductor biomaRt package (v2.40.5)^63^ and were subsequently extracted for the sncRNA-specific analysis, whereas all transcripts were included in the mRNA fragment analysis. For assessment of technical replicates, transcripts were filtered to include at least 2 UMIs in each sample for every compartment.

#### Normalization

Filtered transcripts were normalized using the calcNormFactors() function with the trimmed mean of M values (TMM)-method from the edgeR Bioconductor package (v3.20.9)^64, 65^. For comparison, scaling factors were generated using the relative log expression (RLE) method derived from DESeq2^66^ (Supplementary Fig. 3a). Boxplots of normalized, log2-transformed counts per million (lcmp) values are illustrated in Supplementary Fig. 3b.

#### Detection of differentially expressed transcripts

Differential expression was accessed following a previously reported pipeline (https://f1000research.com/articles/5-1408/v3), which uses linear models. The model was generated using the model.matrix() function from the Limma package (v3.34.9)^67^, which for the first analysis included: disease status (NINDC, INDC, RRMS and SPMS), sex, age and repeated individual, and for the 2^nd^ analysis included RRMS status (Relapse and Remission), sex and age. In the 1^st^ analysis the RRMS vs NINDC (RRMS-NINDC) contrast was explored, whereas for the 2^nd^ analysis the Relapse vs Remission (Relapse-Remission) contrast was applied. Both contrasts were generated using the makeContrasts() function from the Limma package (v3.34.9)^67^. Prior to fitting normalized, lcmp values, the voomWithQualityWeights() function was used to remove heteroscedasity from the data and applying weights based on variance^68, 69^. Processed values were subsequently fitted using the lmFit() function and fitted values were extracted using the fitted() function from Limma (v3.34.9)^67^. The contrasts of interest were applied and differential expression assessed using the contrasts.fit(), eBayes() and topTable() functions from Limma (v3.34.9)^67^.

#### Heatmaps

Normalized, fitted lcmp values from transcripts filtered based on reported (Benjamini-Hochberg (BH) corrected) P-value thresholds were centered and scaled within each compartment using the scale() function of R (v3.4.4) and subsequently visualized using the heatmap.3() function extracted from: https://raw.githubusercontent.com/obigriffith/biostar-tutorials/master/Heatmaps/heatmap.3.R, which requires the gplots (v3.0.1) and devtools (v2.2.1) R packages. Hierarchical clustering within groups (NINDC, SPMS, relapse, remission and INDC) of compartment (PBMCs, CSF cells, Plasma and CSF cells) and transcripts (sub-grouped, if reported) were performed using the hclust(), dist() and as.dendrogram() functions from R (v3.4.4).

### MiRNA target gene prediction and enrichment analyses

Differentially expressed miRNAs were used as input for target prediction using mirDIP (v4.1.11.1). Only predicted target genes with an integrative scores > 0.7 were used for gene ontology (GO) analysis.

The enrichment of predicted target genes of differentially expressed miRNAs among mRNA fragments was calculated using Fisher’s exact test and p < 0.05 was considered significant.

### Gene ontology analyses

GO analysis was performed using Ingenuity Pathway Analysis (IPA, Qiagen) to generate Canonical Pathways. Settings included unbiased parameters for all criteria including tissues selection. Enriched GO terms with adjusted p values < 0.05 (Benjamini-Hochberg) were considered statistically significant.

## Data availability

The RNA-seq data used for sncRNA analyses is available in the Swedish National Data Service (https://snd.gu.se/en, the accession number is pending). Supplementary Data files are available upon request.

## Acknowledgments

This work was supported by the Swedish Research Council, the Swedish Association for Persons with Neurological Disabilities, the Swedish Brain Foundation, the Swedish MS Foundation, the Stockholm County Council (ALF project), AstraZeneca (AstraZeneca-Science for Life Laboratory collaboration), the Margaretha af Ugglas foundation and EU Horizon 2020 (MultipleMS grant, 733161). GZ was supported by a fellowship from the Swedish Society for Medical Research. The authors acknowledge the National Genomics Infrastructure in Stockholm funded by Science for Life Laboratory, the Knut and Alice Wallenberg Foundation and the Swedish Research Council, and SNIC/Uppsala Multidisciplinary Center for Advanced Computational Science for assistance with massively parallel sequencing and access to the UPPMAX computational infrastructure, respectively. The authors also acknowledge input on the initial analysis from Dr. Francesco Marabita and input on writing from Dr. Lara Kular.

## Contributions

MJ, GZ and EP conceived and designed the study. GZ, EP, MHJ and PS conducted experiments. MN, EP, MHJ, GZ performed analysis with the input from DE, OF and MJ. TO, FP, FAN and MK collected and managed samples and clinical information. GZ, EP, MN and MJ wrote the manuscript with assistance from all authors. All authors read and approved the manuscript.

## Competing financial interests

FP has received research grants from Genzyme, Merck KGaA and Novartis, and fees for serving as Chair of DMC in clinical trials with Parexel. TO has received unrestricted MS research grants, honoraria for lectures or advisory boards from Biogen, Novartis, Sanofi, Merck and Roche, none of which has relation to this work. All other authors declare no competing interests.

