## Supplementary Information for "Profiling of small non-coding RNAs across cellular and biofluid compartments: implications for multiple sclerosis immunopathology"

### Supplementary Note 1

In the SPMS group, the profile of the snRNAs resembled RRMS (remission phase) both in PBMCs and in CSF cells (Fig. 3a). In the INDC/SLE group, the snRNAs tended to be upregulated compared to RRMS in PBMCs and downregulated in CSF cells (Fig. 3a).

The majority of snoRNAs demonstrated a tendency for upregulation in SPMS compared to RRMS in PBMCs, while in CSF cells SPMS the snoRNAs profile was more similar to RRMS (Fig. 3b). The snoRNAs displayed a pattern of upregulation (first and second group) and downregulation (third and fourth group) in PBMCs of INDC/SLE compared to RRMS (Fig. 3b). All snoRNAs had a clear tendency of downregulation in INDC/SLE patients compared to RRMS in CSF cells (Fig. 3b).

The profile of the tRNAs in SPMS mimicked the RRMS group profile in PBMCs and the NINDC group profile in CSF cells (Fig. 4). The tRNAs demonstrated a clear pattern of upregulation in PBMCs and downregulation in CSF cells in INDC/SLE patients compared to RRMS (Fig. 4).

In general, the miRNA profile of the SPMS group was more similar to the NINDC profile than to RRMS in all four compartments (Fig. 5a). Due to the complexity of the miRNA patterns, it was difficult to distinguish their profile in the INDC/SLE group.

### Supplementary Figure 1

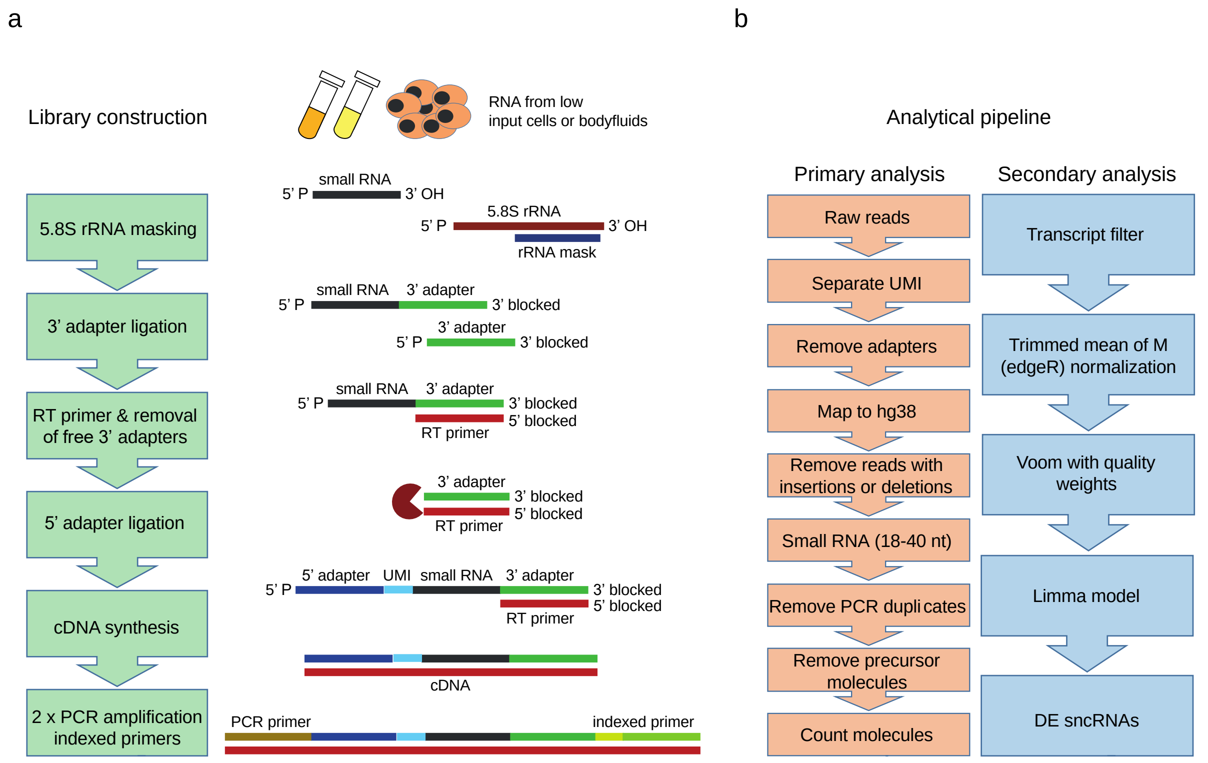


**Supplementary Figure 1. Schematic overview of small non-cording RNA (sncRNA) library preparation and analysis pipeline. (a)** In intracellular RNA, 5.8S ribosomal RNA (rRNA) was masked for adapter ligation immediately by a complementary oligo. The 3' adapter was ligated to RNAs and the free adapter was removed enzymatically prior to the 5' adapter ligation. The 5' adapter comprised a unique molecular identifier (UMI) to remove PCR replicates. Sequences required for Illumina cluster generation and sample indexing were added through two rounds of PCR amplifications. Methodological details are provided in[1, 2]. **(b)** In the primary analysis of the custom analysis pipeline, raw reads were processed and mapped to hg38. Reads with insertions and deletions, as well as reads mapping with less than 18 nt and more than 40 nt, were removed. Based on UMIs, PCR amplicons were collapsed, and subsequently precursor molecules were removed prior to counting the molecules. In the secondary analysis, transcripts were filtered and normalized. Voom with quality weights was applied prior to linear modeling in Limma, which identified differentially expressed sncRNAs within contrasts of interest.

**Supplementary Figure 2**

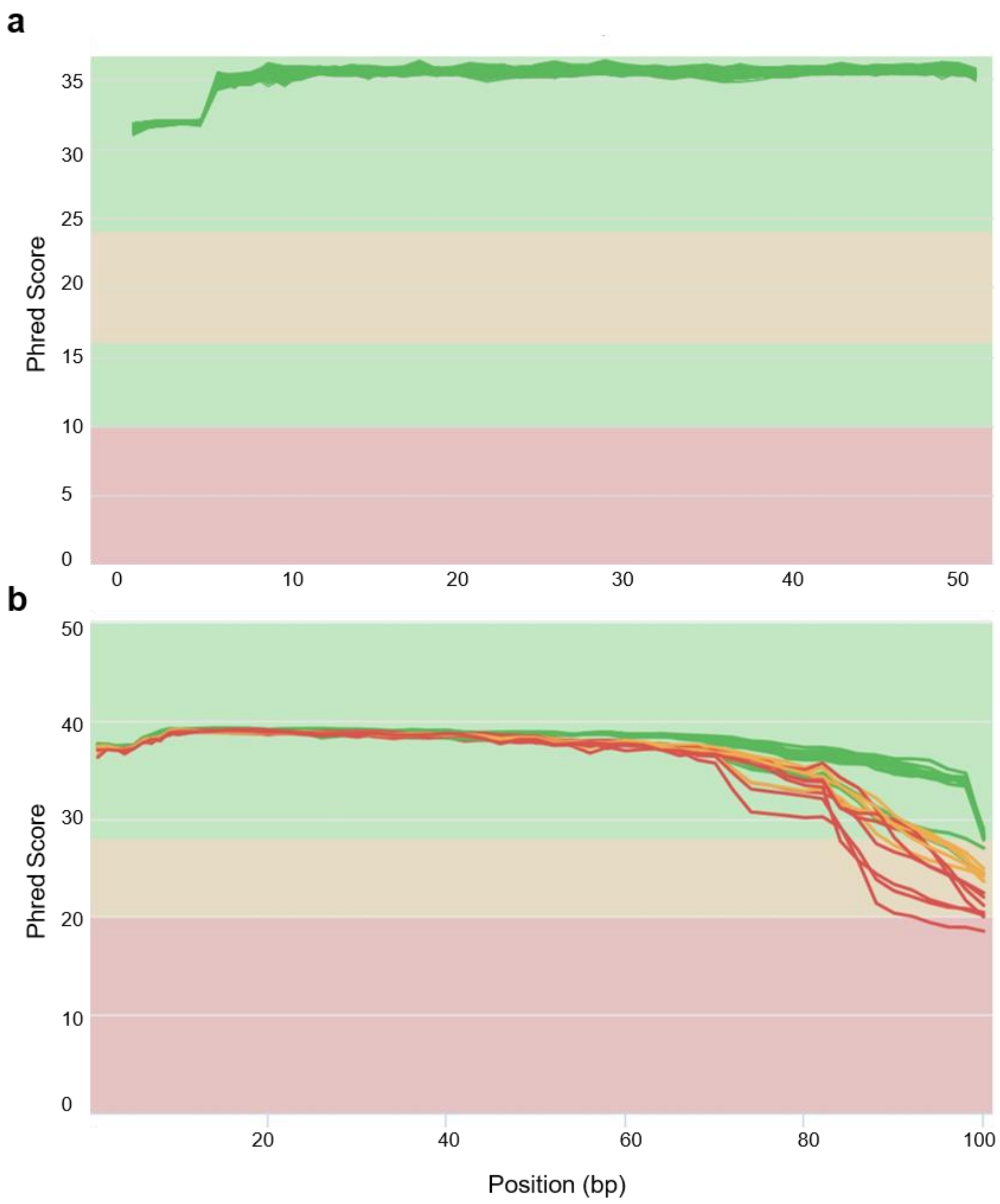


**Supplementary Figure 2. Distribution of quality scores of raw sequence reads of initial (a) and validation sequencing (b) from FastQC software.** MultiQC report showing the mean quality values across each base position in the read. Green, orange and red indicate the samples with Phred quality score above 30, 25 and less than 25 throughout the whole read, respectively.

### Supplementary Figure 3

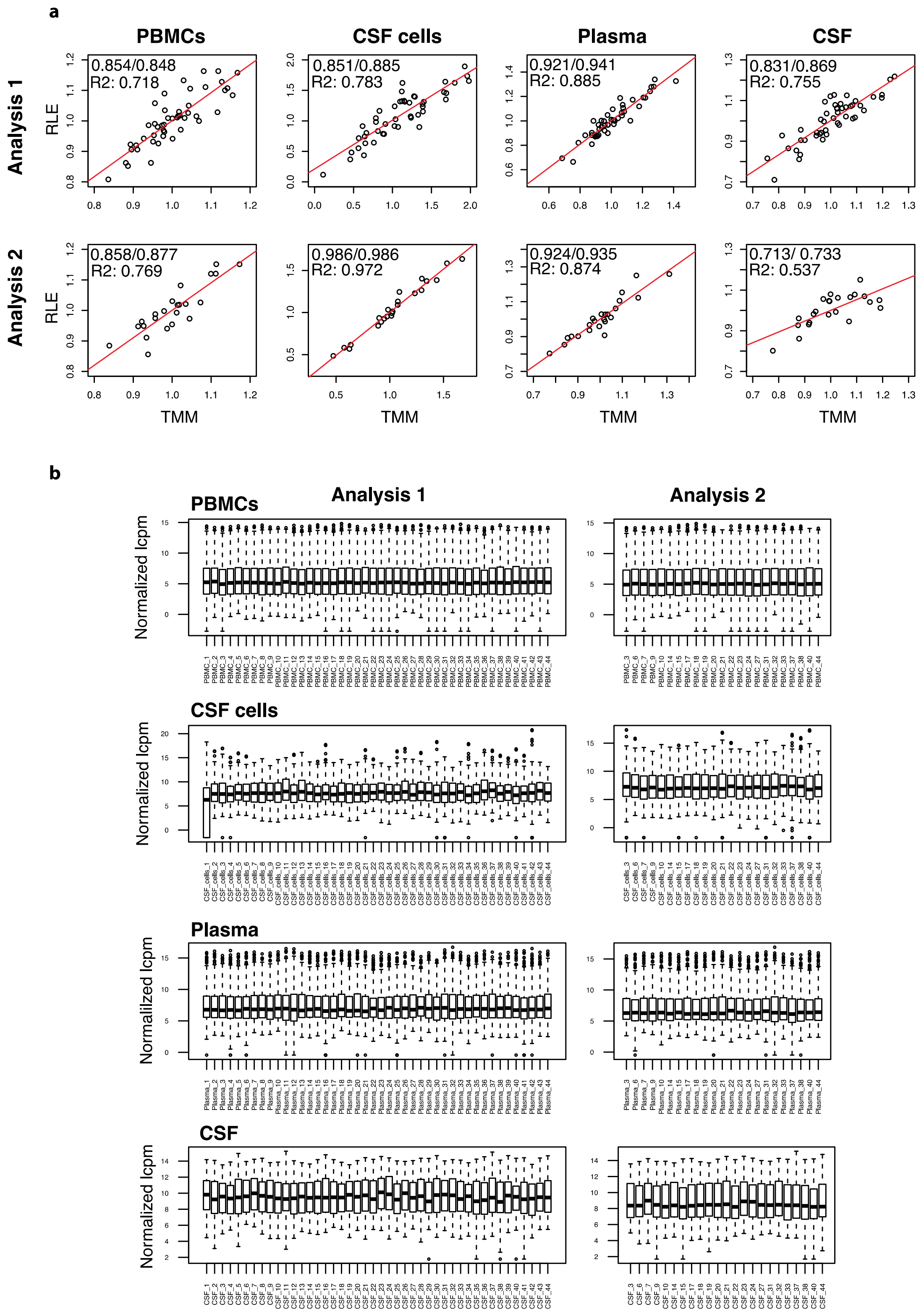

**Supplementary Figure 3. Normalization of expression levels. (a)** Comparison of normalization factors derived from two different methods 1) the trimmed mean of M values (TMM) and 2) relative log expression (RLE), for peripheral blood mononuclear cells (PBMCs), cerebrospinal fluid (CSF) cells, plasma and cell-free CSF compartments, respectively. Analysis 1 focuses on relapsing-remitting multiple sclerosis (RRMS) patients compared to non-inflammatory neurological disease controls (NINDC), whereas analysis 2 focuses on RRMS relapse compared to remission. Spearman/Pearson correlation coefficients as well as R2, are given for each comparison as well as a regression line (red). **(b)** Boxplots of normalized log2 counts per million (lcmp) for PBMCs, CSF cells, plasma and cell-free CSF. The box represents upper and lower quartiles, the black, vertical line represents the median, the whiskers represent maximum and minimum values and circles represent outliers.

**Supplementary Figure 4**

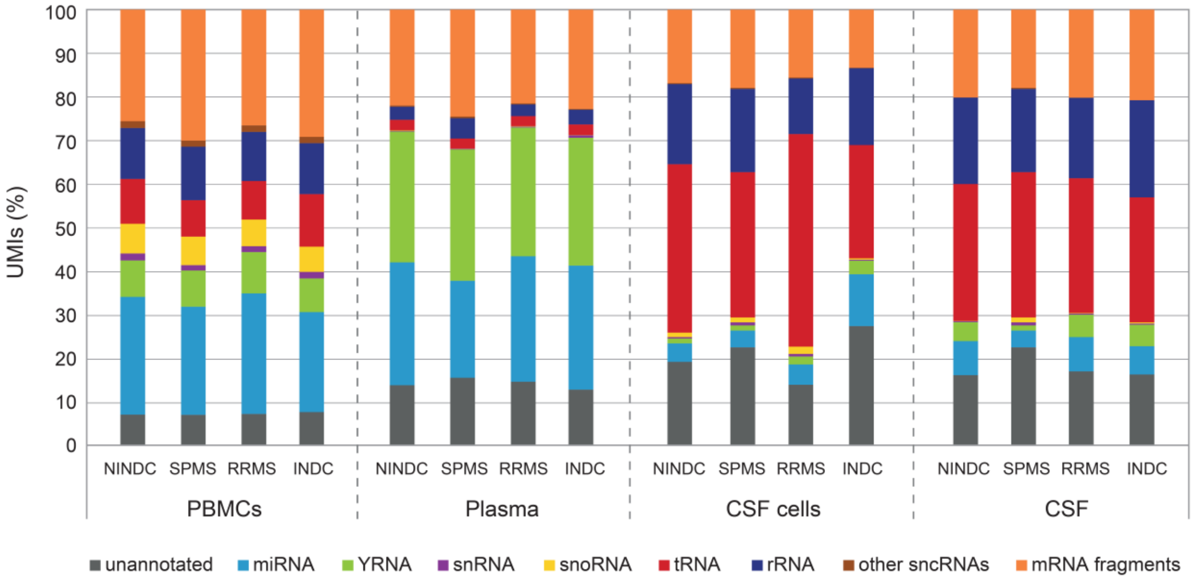


**Supplementary Figure 4. Distribution of small non-coding RNA (sncRNA) fragments across cellular and biofluid compartments and patients' groups.** Unique molecular identifier (UMI) counts across all individuals in the relapsing-remitting multiple sclerosis (RRMS, n=23), secondary progressive multiple sclerosis (SPMS, n=6), non-inflammatory and inflammatory neurological disease controls (NINDC, n=11; INDC, n=5) groups were used to analyze the distribution of different sncRNAs classes in each compartment, including peripheral blood mononuclear cells (PBMCs), plasma, cerebrospinal fluid (CSF) cells and cell-free CSF. Classes of snRNAs include ribosomal RNA (rRNA), microRNA (miRNA), small nuclear RNA (snRNA), small nucleolar RNA (snoRNA), transfer RNA (tRNA), Y RNA (YRNA), sncRNAs in other classes than the aforementioned (other sncRNAs) and messenger RNA (mRNA) fragments. No significant differences in the abundance of RNA classes were observed between different groups of patients (Chi-square test).

### Supplementary Figure 5

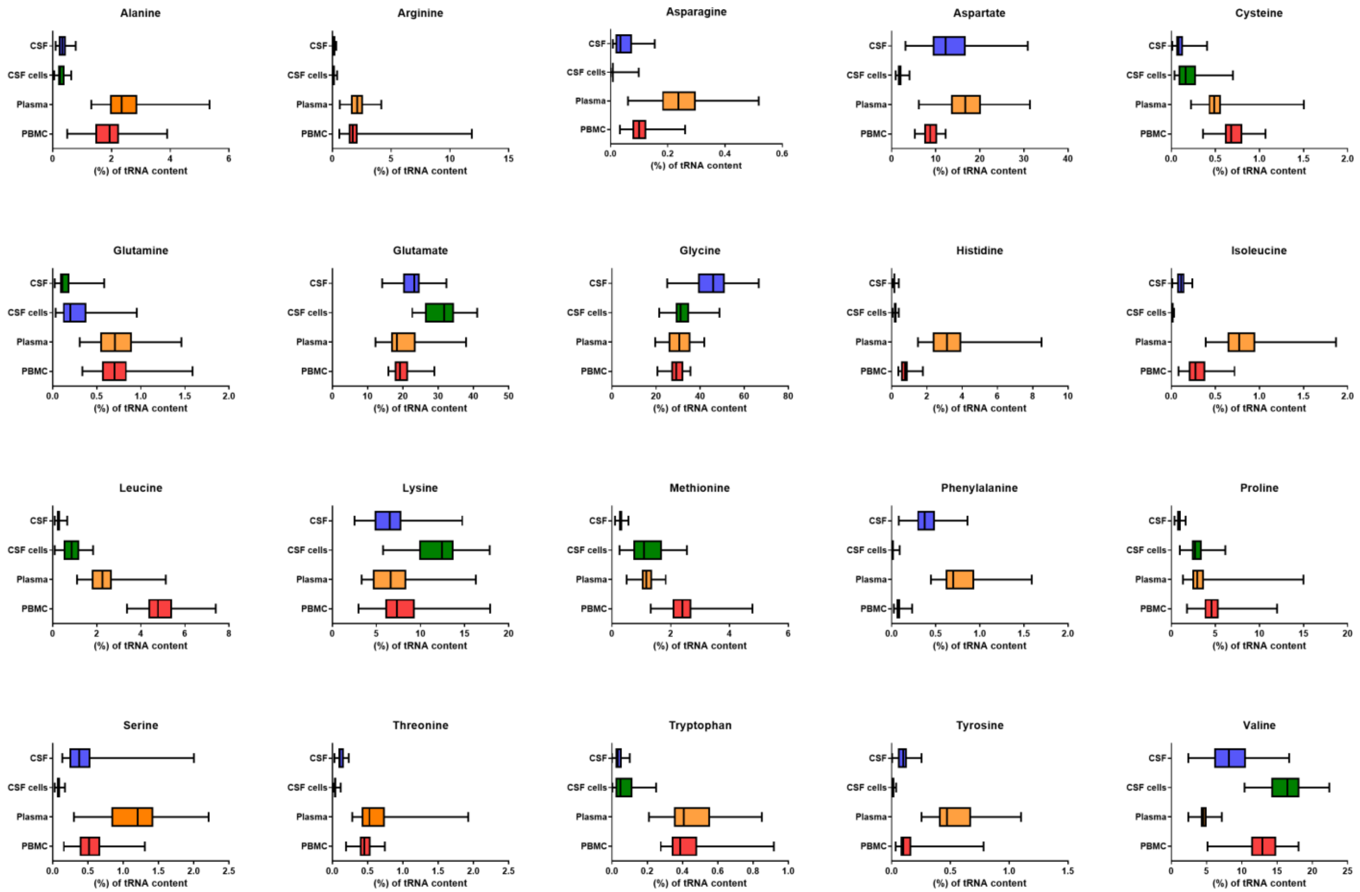


**Supplementary Figure 5. Distribution of transfer RNA (tRNA) fragments for each amino acid across compartments.** All tRNAs for one amino acid were summarized and presented as a fraction of the tRNA counts per compartment, including peripheral blood mononuclear cells (PBMCs), plasma, cerebrospinal fluid (CSF) cells and cell-free CSF.

**Supplementary Table 1.** Sequencing statistics for different compartments.

| | PBMCs | Plasma | CSF cells | CSF |
|---|---|---|---|---|
| **Input reads, summary, M** | 307.5 | 293.5 | 319.9 | 297.1 |
| **Input reads, M, mean (range)** | 6.8 (3.5-11.0) | 6.7 (3.8-10.5) | 7.4 (0.4-31.6) | 6.8 (3.8-10.0) |
| **Uniquely mapped, %, mean (range)** | 27.5 (22.0-34.3) | 15.6 (3.5-61.1) | 7.7 (1.8-35.0) | 10.0 (2.8-51.6) |
| **Mapped to multiple loci, %, mean (range)** | 64.2 (59.1-70.8) | 40.8 (8.6-71.2) | 64.6 (34.7-90.9) | 19.2 (3.8-52.9) |
| **Unmapped, too short, %, mean (range)** | 7.9 (3.2-15.8) | 43.6 (10.1-86.2) | 27.7 (2.5 - 62.0) | 70.7 (34.8-93.4) |
| **Unmapped, other, %, mean (range)** | 0.34 (0.1-0.71) | 0.18 (0.02 -0.60) | 0.05 (0.01- 0.14) | 0.07 (0.01 -0.21) |
| **Unique molecule identifiers (UMIs), M, mean (range)** | 1.8 (1.2-2.6) | 0.38 (0.13-0.92) | 0.67 (0.006-1.82) | 0.09 (0.04-0.21) |

PBMCs – peripheral blood monocular cells, CSF – cerebrospinal fluid, M – million.

**Supplementary Table 2.** Percentage of unique molecular identifier (UMI) counts occupied by the top ten small nuclear RNA (snRNA) fragments.

| PBMC | Plasma | CSF cells |
|---|---|---|
| RNU12 ENST00000362512.1 (15.2%) | RNU5A-1 ENST00000362698.1 (15.0%) | RNU2-2P ENST00000410396.1 (17.9%) |
| RNU2-2P ENST00000410396.1 (13.8%) | RNU12 ENST00000362512.1 (5.2%) | RNU2-36P ENST00000410361.1 (16.5%) |
| RNU5A-1 ENST00000362698.1 (6.5%) | U2 ENST00000619225.1 (2.9%) | RNU5A-1 ENST00000362698.1 (9.5%) |
| RNU2-36P ENST00000410361.1 (6.0%) | RNU2-1 ENST00000618664.1 (2.7%) | RNU12 ENST00000362512.1 (8.3%) |
| U2 ENST00000619225.1 (5.7%) | RNU2-1 ENST00000618602.1 (2.7%) | RNU4-2 ENST00000365668.1 (4.9%) |
| RNU4-2 ENST00000365668.1 (2.3%) | RNU2-1 ENST00000617785.1 (2.7%) | RNU4-1 ENST00000363925.1 (4.7%) |
| RN7SK ENST00000365328.1 (2.1%) | RNU2-1 ENST00000616535.1 (2.6%) | RNU4-46P ENST00000410818.1 (2.8%) |
| RNU5B-1 ENST00000363286.1 (1.5%) | U2 ENST00000613956.1 (2.6%) | U2 ENST00000619225.1 (0.9%) |
| RNU4-1 ENST00000363925.1 (1.4 %) | RNU2-1 ENST00000620268.1 (2.6%) | RNVU1-17 ENST00000384619.1 (0.9%) |
| RNU2-1 ENST00000618664.1 (1.4%) | RNU2-1 ENST00000613778.1 (2.6%) | RN7SK ENST00000365328.1 (0.7%) |

PBMCs – peripheral blood monocular cells, CSF – cerebrospinal fluid.

**Supplementary Table 3.** Percentage of unique molecular identifier (UMI) counts occupied by the top ten small nucleolar RNA (snoRNA) fragments.

| **PBMC** | **CSF cells** |
| --- | --- |
| SNORD104 (8.2%) | SNORD104 (9.9%) |
| SNORD58C (7.2%) | SNORD95 (6.2%) |
| SNORD27 (5.7%) | SNORD27 (4.4%) |
| SNORD95 (4.0%) | SNORD20 (4.6%) |
| SNORD118 (3.2%) | SNORD66 (3.3%) |
| SNORD66 (2.5%) | SNORD100 (2.8%) |
| SNORD65 (2.4%) | SNORD17 (2.4%) |
| SNORD100 (2.2%) | SNORD63 (1.6%) |
| SNORD48 (1.9%) | SNORD26 (1.4%) |
| SNORD17 (1.8%) | SNORD67 (1.3%) |

PBMCs – peripheral blood monocular cells, CSF – cerebrospinal fluid.

**Supplementary Table 4.** Functions and location of altered small nucleolar (snoRNA) and Cajal body-associated (scaRNA) RNA fragments.

| ENST transcript ID | sno/scaRNA | Target RNA/position | Host gene | Group |
|---|---|---|---|---|
| ENST00000391305.1 | SNORA45B | 28S rRNA/U3938 | *RPL27A*, ribosomal protein L27A | 1 |
| ENST00000384581.1 | SNORA61 | 28S rRNA/U2495 | *SNHG12*, small nucleolar RNA host gene 12 | 1 |
| ENST00000384674.1 | SNORA64 | 28S rRNA/U4975 | *RPS2*, ribosomal protein S2 | 1 |
| ENST00000384765.1 | SNORA7A | 28S rRNA/U1569/U1779 | *RPL32*, ribosomal protein L32 | 1 |
| ENST00000384744.1 | SNORA80E | 18S rRNA/U572 and 18S rRNA/U109 | *KHDC4*, KH domain containing 4, pre-mRNA splicing factor | 1 |
| ENST00000408493.2 | SNORA81 | 28S rRNA/U4606 | *EIF4A2*, eukaryotic translation initiation factor 4A2 | 1 |
| ENST00000384714.1 | SNORD15B | 28S rRNA/A3764 | *RPS3*, ribosomal protein S3 | 1 |
| ENST00000364805.1 | SNORD32A | 18S rRNA/G1328 and 28S rRNA/A1511 | *RPL13A*, ribosomal protein L13a | 1 |
| ENST00000625943.1 | SNORD38B | 28S rRNA/A1858 | *RPS8*, ribosomal protein S9 | 1 |
| ENST00000626963.1 | SNORD50B | 28S rRNA/C2848/G2863 | *SNHG5*, small nucleolar RNA host gene 5 | 1 |
| ENST00000581525.1 | SNORD55 | 28S rRNA/C2791 | *RPS8*, ribosomal protein S8 | 1 |
| ENST00000384252.1 | SNORD61 | 18S rRNA/U1442 | *RBMX*, RNA binding motif protein, X-linked | 1 |
| ENST00000391150.1 | SNORD69 | 28S rRNA/G4464 | *GNL3*, guanine nucleotide binding protein-like 3 | 1 |
| ENST00000606577.1 | SNORD96A | 5.8S rRNA/G75 | *GNB2L1*, guanine nucleotide binding protein, component of the 40S ribosomal subunit involved in translational repression | 1 |
| ENST00000516881.1 | SCARNA15 | U2 snRNA/U37 | *SNHG21*, small nucleolar RNA host gene 21 | 2 |
| ENST00000364113.1 | SNORA45A | 28S rRNA/U3899 | *RPL27A*, ribosomal protein L27A | 2 |
| ENST00000364902.1 | SNORA5C | 18S rRNA/U1238 and 18S rRNA/U1625 | *TBRG4*, transforming growth factor beta regulator 4 | 2 |

| | | | | |
|---|---|---|---|---|
| ENST00000408573.1 | SNORD100 | 18S rRNA/G436 | *RPS12*, ribosomal protein S12 | 2 |
| ENST00000408189.1 | SNORD110 | 18S rRNA/U1288 | *NOL5A*, nucleolar protein 5A, a common constituent of C/D box snoRNPs | 2 |
| ENST00000363753.1 | SNORD18A | 28S rRNA/A1313 | *RPL4*, ribosomal protein L4 | 2 |
| ENST00000459623.1 | SNORD19B | 18S rRNA/G683 | *GLN3*, guanine nucleotide binding protein-like 3 | 2 |
| ENST00000363091.1 | SNORD1B | 28S rRNA/G4362 | *SNHG16*, small nucleolar RNA host gene 16 | 2 |
| ENST00000384550.1 | SNORD20 | 18S rRNA/U1804 | *NCL*, nucleolin, involved in the synthesis and maturation of ribosomes | 2 |
| ENST00000383884.1 | SNORD24 | 28S rRNA/C2338/C2352 | *RPL7a*, ribosomal protein L7a | 2 |
| ENST00000365607.2 | SNORD25 | 18S rRNA/G1490 | *SNHG1*, small nucleolar RNA host gene 1, U22 locus | 2 |
| ENST00000363981.1 | SNORD27 | 18S rRNA/A27 | *SNHG1*, small nucleolar RNA host gene 1, U22 locus | 2 |
| ENST00000363961.1 | SNORD36B | 18S rRNA/A668 | *RPL7A*, ribosomal protein L7A | 2 |
| ENST00000516733.1 | SNORD36C | 28S rRNA/A3703 | *RPL7A*, ribosomal protein L7A | 2 |
| ENST00000384048.1 | SNORD37 | 28S rRNA/A3697 | *EEF2*, eukaryotic translation elongation factor 2 | 2 |
| ENST00000365161.1 | SNORD38A | 28S rRNA/A1858 | *RPS8*, ribosomal protein S8 | 2 |
| ENST00000364953.1 | SNORD48 | 28S rRNA/C2279 | *SNHG32*, small nucleolar RNA host gene 32 | 2 |
| ENST00000459174.1 | SNORD4A | 18S rRNA/U121 | *RPL23A*, ribosomal protein L23A | 2 |
| ENST00000626963.1 | SNORD50A | 28S rRNA/C2848 and 28S rRNA/G2863 | *SNHG5*, small nucleolar RNA host gene 5 | 2 |
| ENST00000448188.1 | SNORD57 | 18S rRNA/A99 | *NOL5A*, nucleolar protein 5A, a common constituent of C/D box snoRNPs | 2 |
| ENST00000383875.1 | SNORD58A | 28S rRNA/G4198 | *RPL17*, ribosomal protein L17 | 2 |
| ENST00000607313.1 | SNORD58B | 28S rRNA/G4198 | *RPL17*, ribosomal protein L17 | 2 |

| | | | | |
|---|---|---|---|---|
| ENST00000365223.1 | SNORD58C | 28S rRNA/U4197 and 28S rRNA/G4198 | *RPL17*, ribosomal protein L17 | 2 |
| ENST00000426867.1 | SNORD62B | 18S rRNA/A590 | *PRRC2B*, proline Rich Coiled-Coil 2B, KIAA0515, HLA-B-Associated Transcript 2-Like | 2 |
| ENST00000363214.1 | SNORD68 | 18S rRNA/U428 and 28S rRNA/A2388 | *RPL13*, ribosomal protein L13 | 2 |
| ENST00000365530.1 | SNORD82 | 18S rRNA/A1678 | *NCL*, nucleolin, involved in the synthesis and maturation of ribosomes | 2 |
| ENST00000386745.1 | SNORD83B | unknown, complementary to 18S rRNA | *RPL3*, ribosomal protein L3 | 2 |
| ENST00000584275.1 | SNORD84 | unknown | *BAT1*, ATP-dependent RNA helicase of the DEAD protein family | 2 |
| ENST00000390981.1 | SNORD89 | unknown | *RNF*, ring finger protein 149, ligase activity and ubiquitin-protein transferase activity | 2 |
| ENST00000391145.1 | SNORD90 | unknown | *RC3H2*, ring finger and CCCH-type domains 2 | 2 |
| ENST00000609620.2 | SNORD91A | 28S rRNA/G4588 | *TSR1*, ribosome maturation factor | 2 |
| ENST00000608459.2 | SNORD91B | 28S rRNA/G4588 | *TSR1*, ribosome maturation factor | 2 |
| ENST00000408813.1 | SNORD93 | 18S rRNA/A576 | *SNHG26*, small nucleolar RNA host gene 26 | 2 |
| ENST00000408612.1 | SNORD99 | 28S rRNA/A2774 | *SNHG12*, small nucleolar RNA host gene 12 | 2 |
| ENST00000364938.1 | SNORA73A | unknown | *RCC1*, regulator of chromosome condensation | 3 |
| ENST00000363217.1 | SNORA73B | unknown | *RCC1*, regulator of chromosome condensation | 3 |
| ENST00000362883.1 | SNORD104 | 28S rRNA/C1327 | *SNHG25*, small nucleolar RNA host gene 25 | 3 |
| ENST00000364009.1 | SNORD14E | unknown | *Hsp70*, heat shock protein family A member 8 | 3 |
| ENST00000390930.1 | SNORD17 | 28S rRNA/U3797 | *SNX5*, sorting nexin 5 | 3 |
| ENST00000383953.1 | SNORD21 | 28S rRNA/G1303 | *RPL5*, ribosomal protein L5 | 3 |

| | | | | |
|---|---|---|---|---|
| ENST00000384147.1 | SNORD26 | 28S rRNA/A389 | *SNHG1*, small nucleolar RNA host gene 1, U22 locus | 3 |
| ENST00000384693.1 | SNORD30 | 28S rRNA/A3804 | *SNHG1*, small nucleolar RNA host gene 1, U22 locus | 3 |
| ENST00000362761.1 | SNORD33 | 18S rRNA/U1326 | *RPL13A*, ribosomal protein L13a | 3 |
| ENST00000365444.1 | SNORD6 | 28S rRNA/G2411 | *TAF1D*, TATA-box binding protein associated factor, RNA polymerase I subunit D | 3 |
| ENST00000428514.1 | SNORD62A | 18S rRNA/A590 | *PRRC2B*, proline Rich Coiled-Coil 2B, HLA-B-Associated Transcript 2-Like | 3 |
| ENST00000384262.1 | SNORD63 | 28S rRNA/A4541 | *HSPA9B*, heat Shock Protein Family A Member 9 | 3 |
| ENST00000390856.1 | SNORD66 | 18S rRNA/C1272 | *EIF4G1*, eukaryotic translation initiation factor 4 gamma 1 | 3 |
| ENST00000390833.1 | SNORD67 | U6 snRNA/C60 | *CKAP5*, cytoskeleton associated protein 5 | 3 |
| ENST00000411292.1 | SNORD71 | 5.8S rRNA/U14 | *AP1G1*, adaptor-related protein complex 1, gamma 1 subunit | 3 |
| ENST00000386747.1 | SNORD83A | unknown | *RPL3*, ribosomal protein L3 | 3 |
| ENST00000579879.1 | SNORD95 | 28S rRNA A2802/C2811 | *GNB2L1*, guanine nucleotide binding protein, component of the 40S ribosomal subunit involved in translational repression | 3 |
| ENST00000515982.1 | SCARNA6 | U5 snRNA/U41 | *ATG16L1*, autophagy related 16 like 1 | 4 |
| ENST00000516989.1 | SCARNA6 | U5 snRNA/U41 | *ATG16L1*, autophagy related 16 like 1 | 4 |
| ENST00000408876.1 | SNORD23 | unknown | *NOP53*, ribosome biogenesis factor | 4 |

**Supplementary Table 5.** Percentage of unique molecular identifier (UMI) counts occupied by the top ten transfer RNA (trna) fragments.

| **PBMC** | **Plasma** | **CSF cells** | **CSF** |
|---|---|---|---|
| trna5-GlyGCC_1 (3.6%) | trna5-GlyGCC_1 (5.1%) | trna87-GluCTC_1 (3.6%) | trna5-GlyGCC_1 (4.3%) |
| trna59-GluCTC_1 (3.5%) | trna4-AspGTC_1 (3.1%) | trna77-GluCTC_2 (3.6%) | trna68-GlyGCC_1 (4.1%) |
| trna19-GlyGCC_2 (2.5%) | trna19-GlyGCC_2 (2.8%) | trna71-GluCTC_1 (3.6%) | trna128-GlyGCC_1(4.1%) |
| trna128-GlyGCC_1 (2.4%) | trna77-GluCTC_1 (2.6%) | trna80-GluCTC_1 (3.6%) | trna24-GlyGCC_1 (4.0%) |
| trna68-GlyGCC_1 (2.4%) | trna48-AspGTC_1 (2.6%) | trna74-GluCTC_1 (3.6%) | trna19-GlyGCC_1 (4.0%) |
| trna24-GlyGCC_1 (2.4%) | trna45-AspGTC_1 (2.3%) | trna77-GluCTC_1 (3.5%) | trna19-GlyGCC_2 (4.0%) |
| trna19-GlyGCC_1 (2.3%) | trna87-GluCTC_1 (2.1%) | trna59-GluCTC_1 (3.4%) | trna25-GlyGCC_1 (4.0%) |
| trna25-GlyGCC_1 (2.3%) | trna24-GlyGCC_1 (2.1%) | trna68-GlyGCC_1 (2.7%) | trna133-GlyCCC_1 (3.9%) |
| trna77-GluCTC_1 (2.3%) | trna77-GluCTC_2 (2.1%) | trna19-GlyGCC_1 (2.7%) | trna18-GlyGCC_1 (3.9%) |
| trna18-GlyGCC_1 (2.3%) | trna80-GluCTC_1 (2.1%) | trna24-GlyGCC_1 (2.7%) | trna4-GlyCCC_1 (3.9%) |

PBMCs – peripheral blood monocular cells, CSF – cerebrospinal fluid.

**Supplementary Table 6.** Percentage of unique molecular identifier (UMI) counts occupied by the top ten microRNAs.

| PBMC | Plasma | CSF cells | CSF |
|---|---|---|---|
| miR-26a (7.2%) | miR-486-5p (11.7%) | miR-26a-5p (12.8%) | miR-99a-5p (11.7%) |
| miR-92a (5.6%) | miR-26a-5p (9.4%) | miR-92a-3p (7.2%) | miR-26a-5p (11.3%) |
| miR-21 (3.9%) | miR-451a (6.4%) | miR-486-5p (4.6%) | miR-143-3p (9.2%) |
| let-7f (3.7%) | miR-92a-3p (6.4%) | let-7f-5p (4.4%) | miR-204-5p (5.1%) |
| let-7a (3.4%) | miR-144-3p (3.6%) | miR-150-5p (4.1%) | miR-125b-5p (4.9%) |
| miR-191 (3.4%) | miR-199a-5p (3.3%) | miR-451a (4.0%) | miR-125a-5p (4.2%) |
| miR-181a (2.6%) | miR-16-5p (3.2%) | let-7a-5p (3.8%) | miR-100-5p (3.9%) |
| miR-103 (2.3%) | miR-191-5p (2.6%) | miR-146a-5p (3.2%) | miR-30a-5p (2.8%) |
| miR-30d (2.3%) | miR-142-5p (2.4%) | miR-142-5p (2.8%) | miR-10a-5p (2.7%) |
| let-7i (2.0%) | miR-30d-5p (2.4%) | miR-21-5p (2.6%) | miR-124-3p (2.0%) |

PBMCs – peripheral blood monocular cells, CSF – cerebrospinal fluid.

**Supplementary Table 7.** Overlap of predicted target genes of differentially expressed microRNAs (miRNAs) with mRNA fragments.

| Comparison | DE miRNA | p-value |
|---|---|---|
| **RRMS vs NINDC** | | |
| *PBMC* | miR-548o-3p | 0.12 |
| *CSF cells* | miR-146a-5p | 0.14 |
| | miR-148a-3p | 0.07 |
| | miR-181a-5p | 0.64 |
| | miR-181b-5p | 0.64 |
| | miR-204-5p | 0.79 |
| | miR-29a-3p | 0.78 |
| | miR-29b-3p | 0.78 |
| | miR-30a-5p | 0.62 |
| | miR-30c-5p | 0.62 |
| | miR-342-3p | 0.93 |
| ***Relapse vs Remission*** | | |
| *CSF cells* | let-7a-5p | **0.03** |
| | let-7d-5p | **0.03** |
| | let-7d-3p | N.A. |
| | let-7f-5p | **0.02** |
| | let-7i-5p | **0.02** |
| | miR-125a-5p | **0.04** |
| | miR-140-3p | **0.006** |
| | miR-142-5p | **0.002** |
| *CSF cells cont.* | miR-146a-5p | **0.0004** |
| | miR-146b-5p | **0.0007** |
| | miR-148a-3p | **0.03** |
| | miR-150-5p | **0.04** |
| | miR-151a-3p | **0.02** |
| | miR-155-5p | **0.005** |
| | miR-181a-5p | **0.049** |
| | miR-186-5p | 0.057 |
| | miR-191-5p | **0.03** |
| | miR-192-5p | **0.04** |
| | miR-21-3p | **0.007** |
| | miR-25-3p | **0.02** |
| | miR-26a-5p | **0.02** |
| | miR-30d-5p | **0.004** |
| | miR-30e-3p | **0.02** |
| | miR-320a | 0.14 |
| | miR-3609 | **0.001** |
| | miR-362-5p | **0.02** |
| | miR-363-3p | **0.01** |
| | miR-425-5p | **0.01** |
| | miR-451a | **0.01** |
| | miR-486-5p | **0.02** |
| | miR-532-5p | **0.002** |
| | miR-92a-3p | **0.01** |
| | miR-93-5p | **0.03** |

Enrichment of predicted target genes of differentially expressed microRNAs (miRNAs) with mRNA fragments differentially expressed in peripheral blood mononuclear cells (PBMCs) and cerebrospinal fluid (CSF) cells between relapsing-remitting multiple sclerosis (RRMS) patients compared to non-inflammatory other neurological disease controls (NINDC) as well as RRMS relapse compared to remission was calculated using the Fisher's exact test. DE miRNAs - differentially expressed miRNAs, p-value – significant in bold, N.A. – no predicted targets available on *mirDIP* with 'Very High' minimum score.
